# Simulation-guided engineering of antibiotics for improved bacterial uptake

**DOI:** 10.1101/2020.10.08.330332

**Authors:** Ricardo J. Ferreira, Valeria Aguilar, Ana M. Villamil Giraldo, Peter M. Kasson

**Affiliations:** Science for Life Laboratory, Department of Cell and Molecular Biology, Uppsala University, BMC Box 596, S-751 23 Uppsala, Sweden; Departments of Biomedical Engineering and Molecular Physiology and Biological Physics, University of Virginia, Charlottesville, VA 22908, USA

## Abstract

The Gram-negative bacterial outer membrane poses a major obstacle to the development of much-needed antibiotics against drug-resistant infections. Its chemical composition and porin proteins differ from Gram-positive bacteria and mammalian cells, and heuristics developed for mammalian cell uptake apply poorly. Recently, machinelearning methods have predicted small-molecule uptake into Gram-negative bacteria, offering the possibility to rationally optimize this aspect of antibiotic lead development. Here, we report physics-based methods to prospectively predict Gram-negative bacterial uptake, select, and synthesize promising chemical derivatives targeting *E. coli* DNA gyrase B. Our methods do not require empirical parameterization and are readily adaptable to new chemical scaffolds. These physics-based predictions well capture experimentally measured uptake (r > 0.95) and are indeed predictive of antimicrobial activity (r > 0.92). These methods can be used prospectively in combination with target-binding simulations to optimize both bacterial uptake and target binding, overcoming important barriers to antibiotic lead generation before small-molecule synthesis.

## Introduction

Antibiotic-resistant bacterial infections are an increasing global health threat and are estimated to surpass cancer as a cause of death by 2050^1^. Responding to this challenge requires improved diagnostics, improved antimicrobial stewardship, and critically the development of new antibiotics^2, 3, 4^. New classes of antibiotics active against Gramnegative bacteria have been particularly difficult to develop because in order to reach their targets antibiotics must traverse the Gram-negative outer membrane, which poses a permeability barrier^5, 6, 7^ Depending on the antibiotic chemistry, inner membrane permeability and efflux can also be substantial contributors, but here we concentrate on the outer membrane, which is a major determining factor for several major classes of antibiotics. This outer membrane differs from both mammalian cells and gram-positive bacterial membranes in the high proportion of lipopolysaccharide (LPS) in its outer leaflet and the semi-specific uptake porins, creating very different criteria for efficient antibiotic uptake^8, 9, 10^. In fact, most FDA-approved antibiotics against Gram-negative bacteria violate at least one of the heuristic rules developed to predict mammalian cell uptake^5^. Approaches for selecting and optimizing lead compounds largely rely upon empirical measurements of bacterial growth inhibition, and it has been much easier to rationally or computationally optimize on-target activity than cellular accumulation.

Computational approaches to optimize compound accumulation in Gramnegative bacteria have used a combination of machine learning on molecular features and physics-based methods. Gram-negative accumulation results from a combination of uptake, efflux, and intracellular metabolism. For many hydrophilic compounds, uptake is a good predictor of intracellular accumulation^11, 12, 13^. Hergenrother and colleagues have reported good success with machine learning based on accumulation data^14, 15^. An alternate approach has been to parameterize molecular descriptors based on physics-based simulations^16, 17, 18, 19, 20, 21^ or to forgo computational optimization and focus solely on medium-scale screening approaches^22, 23, 24^. We have previously shown how a simulation-based approach can prospectively predict antibiotic uptake without explicit parameterization^12^. Here, we test the ability of such an approach to guide compound selection for increased uptake and increased antimicrobial potency prior to chemical synthesis.

DNA gyrase B has been an attractive yet challenging target for antibacterial drug development^25, 26, 27, 28, 29, 30^. The most successful inhibitor of this target, novobiocin, was withdrawn due to lack of safety or effectiveness^31^. The ATP-binding pocket of DNA gyrase B has been the subject of renewed preclinical interest, but Gram-negative uptake has been a persistent challenge^30^. This is evidenced by poor minimum inhibitory concentrations (MICs) even when biochemical inhibitory concentrations (IC_50_s) against *E. coli* DNA gyrase B are reported in the nanomolar range. As a proof of concept, we selected flavanone compounds for optimization. Flavonoids have been the subject of previous medicinal chemistry efforts against E. coli DNA gyrase B^32, 33, 34^, and the flavanone naringenin has been a particularly promising starting point, as it provides a readily derivatized scaffold.

In this paper we aim to optimize the bacterial permeabilities for a small set of naringenin hydrazone derivatives while maintaining or improving binding affinity to *E. coli* DNA gyrase B. Prior to synthesizing these derivatives, we estimated both the relative free-energies of binding and the outer-membrane permeability, using calculated OmpF permeability as a surrogate for full E. coli outer membrane permeability as we have previously demonstrated^12^. Improved outer-membrane permeability and equal or better binding affinity were used as selection criteria for synthesizing derivatives and testing them experimentally. The resulting experimental data on both outer-membrane uptake and cellular growth inhibition correlate well with our predictions, demonstrating that molecular simulations of membrane permeability can be used in a prospective manner to guide selection and chemical synthesis of potential antibacterial lead compounds.

## Results

To test the hypothesis that outer-membrane permeability simulations can guide antibiotic lead optimization, we selected a set of synthetically accessible naringenin derivatives for simulation and assessment. The naringenin scaffold can bind and inhibit the ATP-binding pocket of *E. coli* DNA gyrase B with moderate affinity and, we predict, poor-to-moderate permeability, leading to a suboptimal MIC in bacteria (428 μg/mL). Naringenin is commercially available and readily derivatized, providing a good proof-of-concept system for optimization of antibiotic leads based on prospective prediction of bacterial uptake. Hydrazone derivatives were selected based on examination of the binding pose of naringenin to DNA gyrase B and the potential to reinforce hydrogen bonding with Asn46 and Asp73 (see Supplement for further discussion).

We therefore predicted the *E. coli* outer-membrane permeability (here estimated as permeability via OmpF, see prior studies^12, 16^ for discussion of this approximation) and the approximate target binding affinity of a set of 29 derivatives, as described below. Predicted permeability and approximate target binding are listed in Table S2. 61 derivatives (hydrazone-linked flavanones, see Methods) were initially evaluated by molecular docking. Of these, 29 were predicted to maintain a binding mode similar to that of a previously co-crystallized inhibitor (Figure 1A; Tables S1-S2 and Figures S1-S2). Approximate receptor-binding energies and membrane permeabilities were predicted for each of these. Six of these compounds (21%) were predicted to have lower permeabilities than naringenin (Figure 1B). Herein, compounds D19 (carboxylate derivative) D29 (quinoxaline), D5 (*m*-ethyl benzoate), D13 (*o*-nitrophenyl), D15 (indan), and D21 (imidazole) were predicted to have permeabilities lower than 1 × 10^-8^ cm/s. Approximate binding energies calculated via MM-PBSA are listed in the Supplement. These compounds were then filtered based on 1) prediction of improved bacterial permeability, 2) prediction of maintained or improved target binding, and 3) commercial availability of starting materials to yield the compound set shown in Figure 2. Compound **1** is the starting compound; compounds **2-4** are previously synthesized compounds^35^ that were also predicted to have binding energies and permeabilities within the filter criteria; compounds **5-9** were synthesized based on the criteria above. Compound **10** was synthesized as a negative control based on its poor predicted permeability.

**Figure 1.**
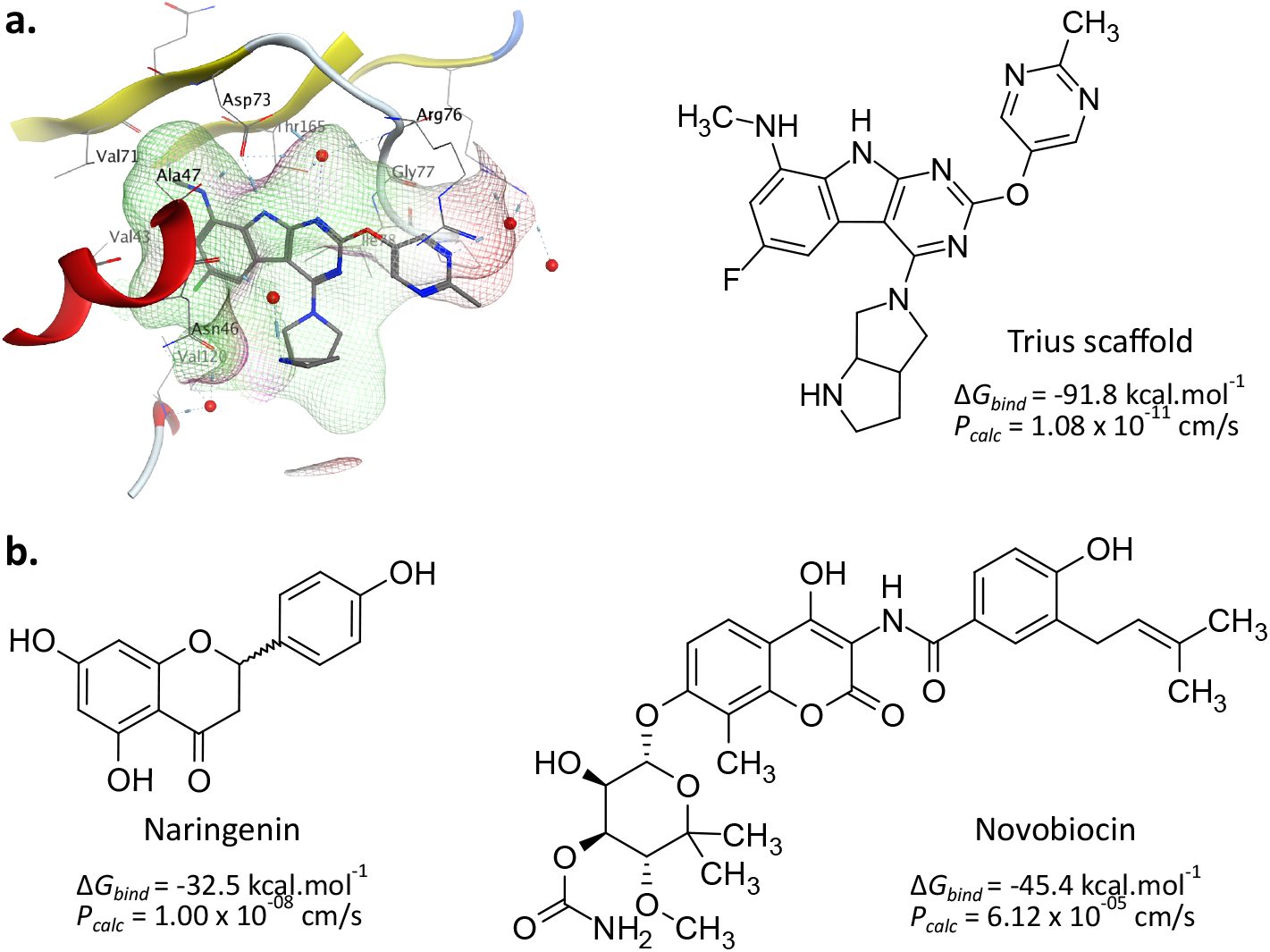
DNA gyrase B binding pocket and inhibitors. Rendered in panel (a) is the ATP-binding pocket of *E. coli* DNA gyrase B with an inhibitor bound, based on co-crystal structure 4KFG. The chemical structure of the co-crystalized ligand is shown with approximate binding free energy (MM-PBSA) and predicted outer-membrane permeability. Panel (b) similarly displays the naringenin starting scaffold and the related antibiotic novobiocin with their predicted binding and permeability values.

**Figure 2.**
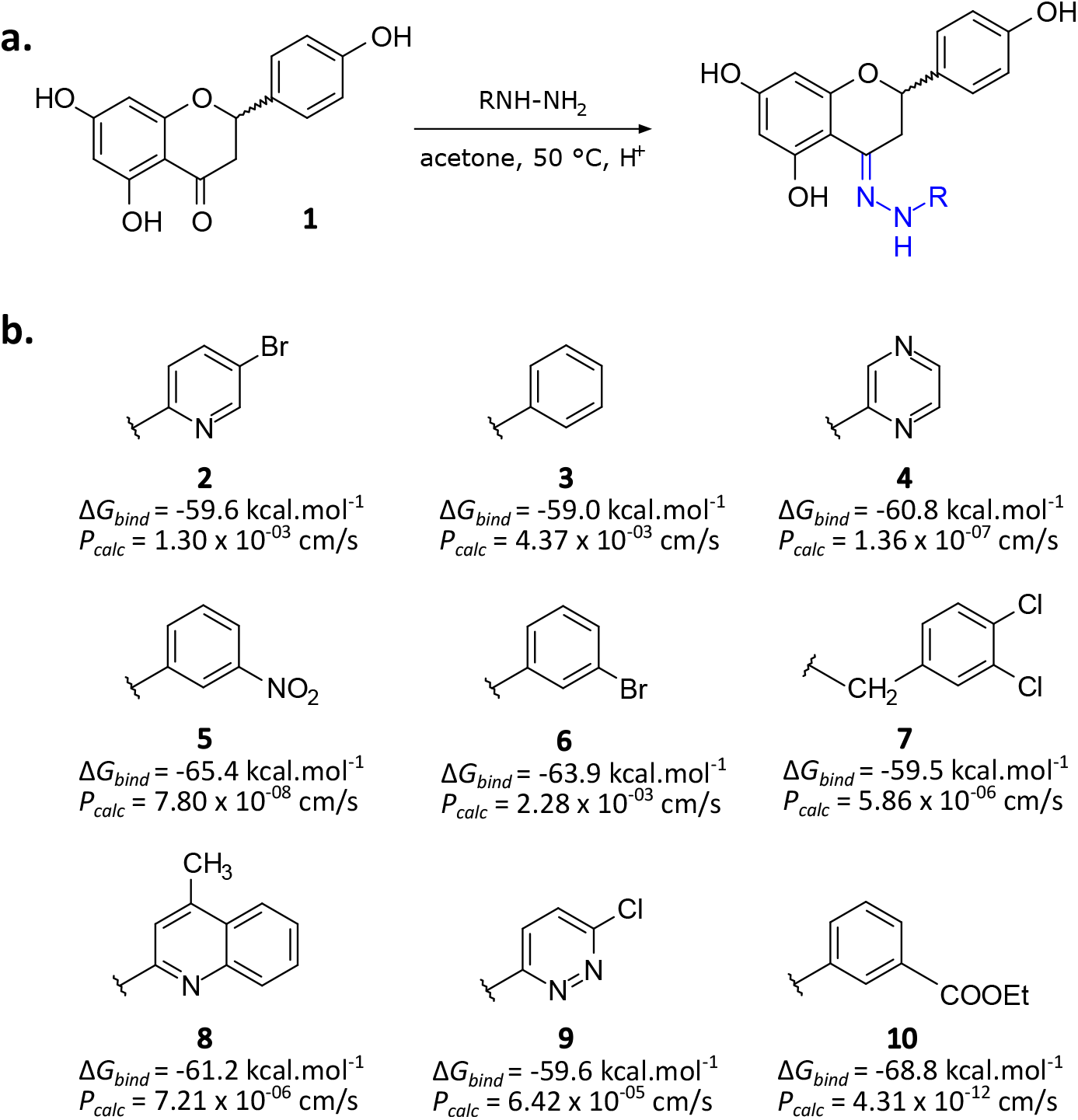
Naringenin derivatives tested. Schematized in panel (a) is the derivatization strategy used to prepare derivatives of naringenin (**1**). Reaction conditions were hydrazine (R-NH-NH_2_) (2.0 eq.) in ethanol with a catalytic amount of acetic acid, in a reflux apparatus for 12-48 h. Schematized in panel (b) are the naringenin derivatives predicted and tested, with their predicted approximate binding free energies and permeabilities. All biological activities and physicochemical properties can be found in Tables 1 and S3).

Selected compounds were synthesized and tested experimentally for bacterial outer membrane uptake, inhibition of DNA gyrase, and inhibition of bacterial growth. The compound set also included three previously synthesized flavanone derivatives and three comparator compounds predicted to have low bacterial membrane permeability. Synthesis and characterization is described in the Methods. Bacterial outer membrane uptake was measured using an assay for osmotic swelling of bacterial outer membrane vesicles (OMV) that we have reported previously^12^. Results of this assay show a strong correlation (Pearson *r* 0.95) of outer-membrane uptake of the synthesized derivatives with predicted permeabilities over an 8-log range (Fig. 3). Notably, compounds **2** (*P_calc_* 1.30 × 10^-3^ cm/s; *P_exp_* 99.6%), **3** (*P_calc_* 4.37 × 10^-3^ cm/s; *P_exp_* 103.8%) and **6** (*P_calc_* 2.28 × 10^-3^ cm/s; *P_exp_* 85.2%) had permeabilities comparable to that of glycine (*P_calc_* 5.32 × 10^-2^ cm/s; *P_exp_* 100%). Our prior validation studies on simulated uptake via OmpF and OMV swelling show similarly good correlations between simulations and accumulation measured via mass spectrometry^12^ and also present validation data on OmpF-deletion strains. The logarithmic relationship between permeability coefficients and osmotic swelling is well described^36^, and the final concentration of antibiotic scales with OMV swelling.

**Figure 3.**
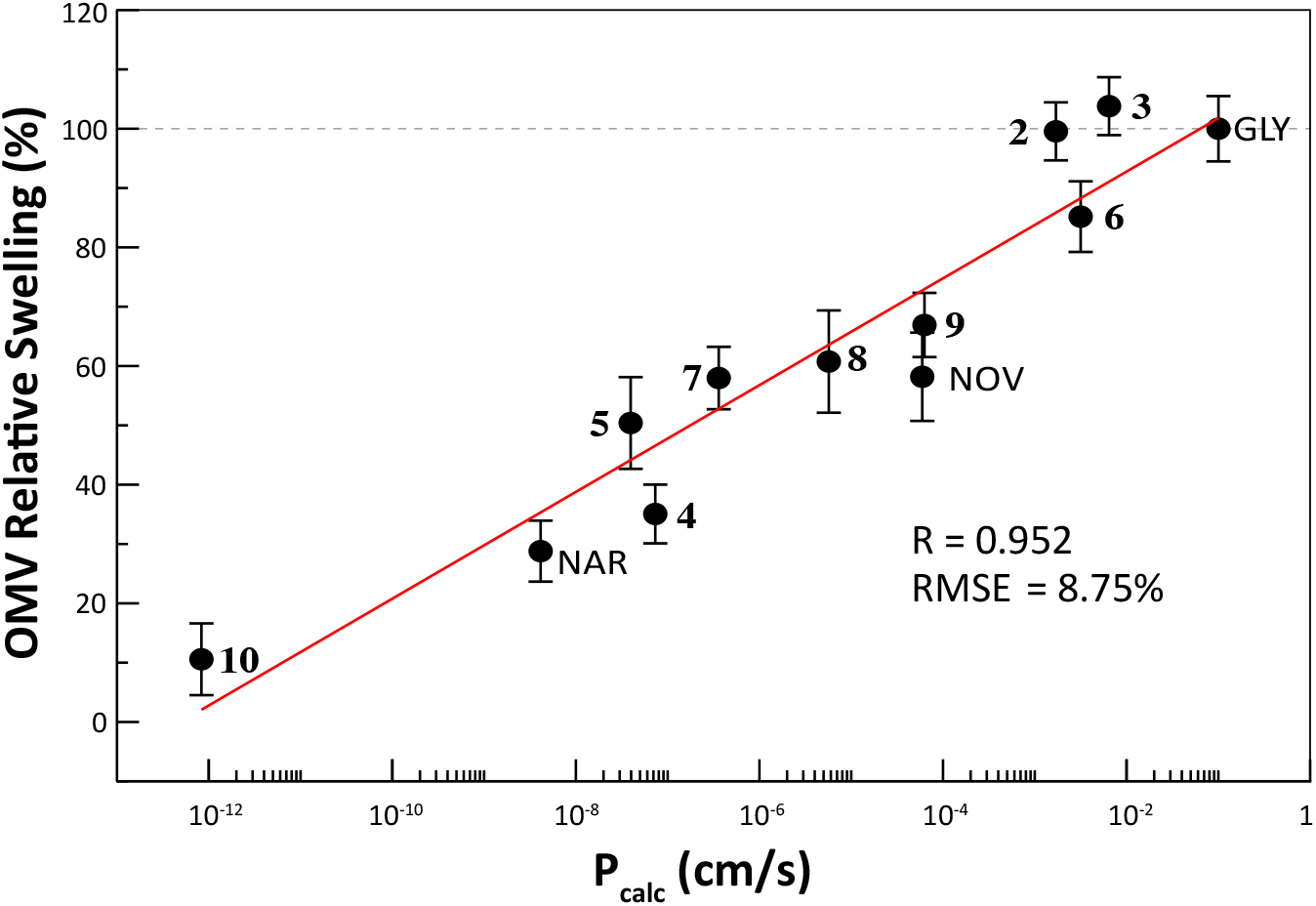
Predicted permeabilities correlate well with measured outer-membrane swelling. Plotted are semilogarithmic correlations of experimental OMV swelling rates and predicted permeabilities in *E. coli* outer membrane vesicles containing OmpF for naringenin (NAR), glycine (GLY), novobiocin (NOV) and the flavanone derivatives **2-10**. Permeabilities are normalized with respect to glycine.

We next tested the antibacterial activity of each compound against the *E. coli* strain MG1655. Minimum inhibitory concentrations (MIC) were determined via microbroth dilution and EC_50_ values were plotted against predicted outer-membrane permeability values in Fig. 4. Again, EC_50_ values show a strong correlation with the logarithm of predicted permeability (Pearson *r* 0.93), with 86% of variance in MIC explained by predicted permeability. Compounds **6, 7** and **10** also displayed a moderate bacteriostatic effect at lower concentrations, retaining 20 to 40% growth inhibition as low as 1.0 μg/mL. EC_50_ for ciprofloxacin and novobiocin were measured as controls at 4 ng/mL and 7.9 μg/mL, respectively. The strong correlation suggests that membrane permeability is a first-order determinant of antimicrobial activity for the compounds tested, in accordance with previous predictions^12^. These data take the additional step to demonstrate that permeability can be optimized prospectively based on simulations to improve antimicrobial activity.

**Figure 4.**
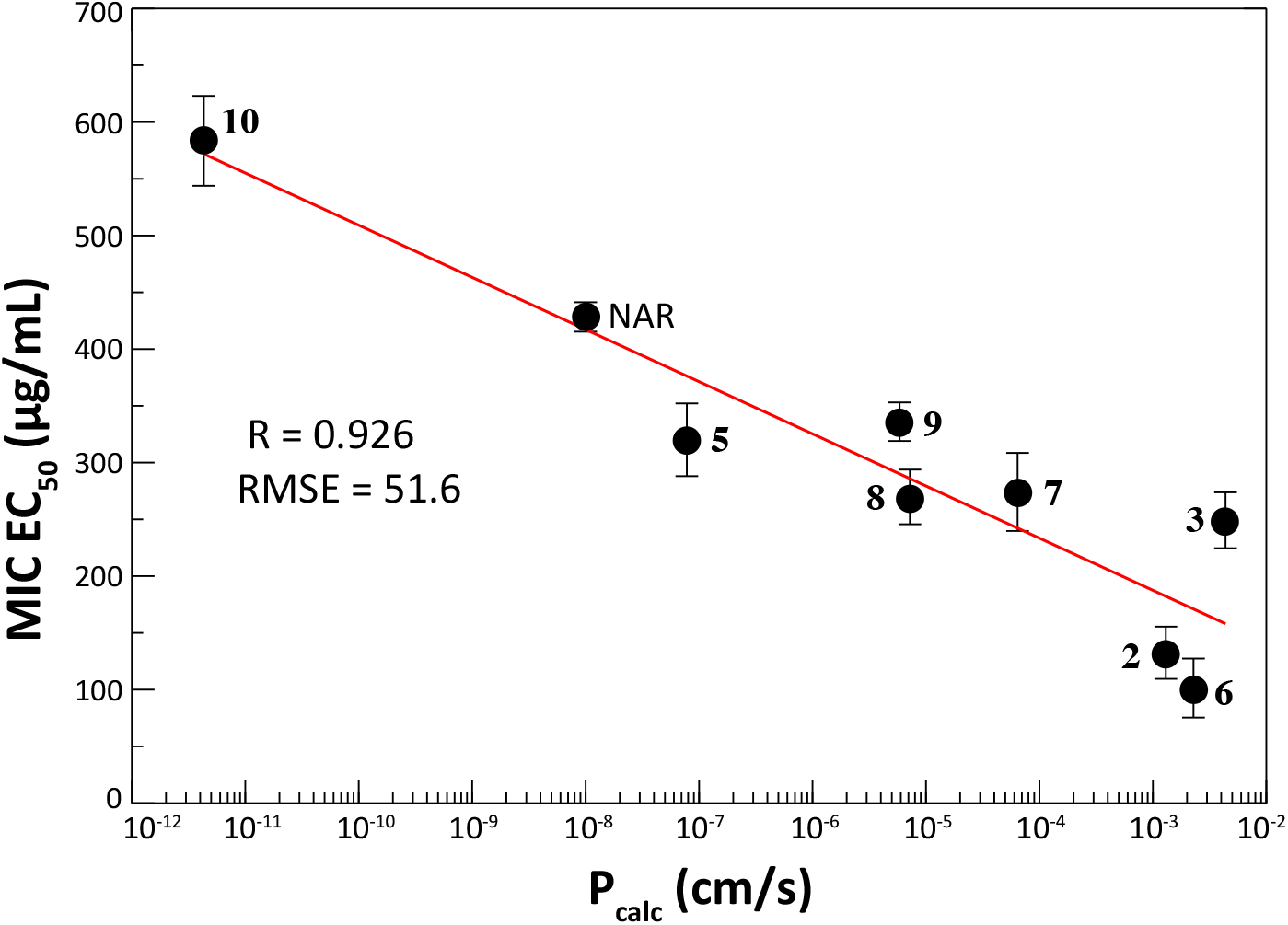
Predicted outer-membrane permeabilities correlate well with antimicrobial activity. Plotted are semilogarithmic correlations of the growth inhibitory concentrations (MIC EC_50_) with calculated OmpF outer-membrane permeabilities (*P_calc_*) for naringenin (NAR) and all flavanone derivatives except **4**. Compound **4** EC_50_ was determined to be > 400 μg/mL.

**Table 1.**
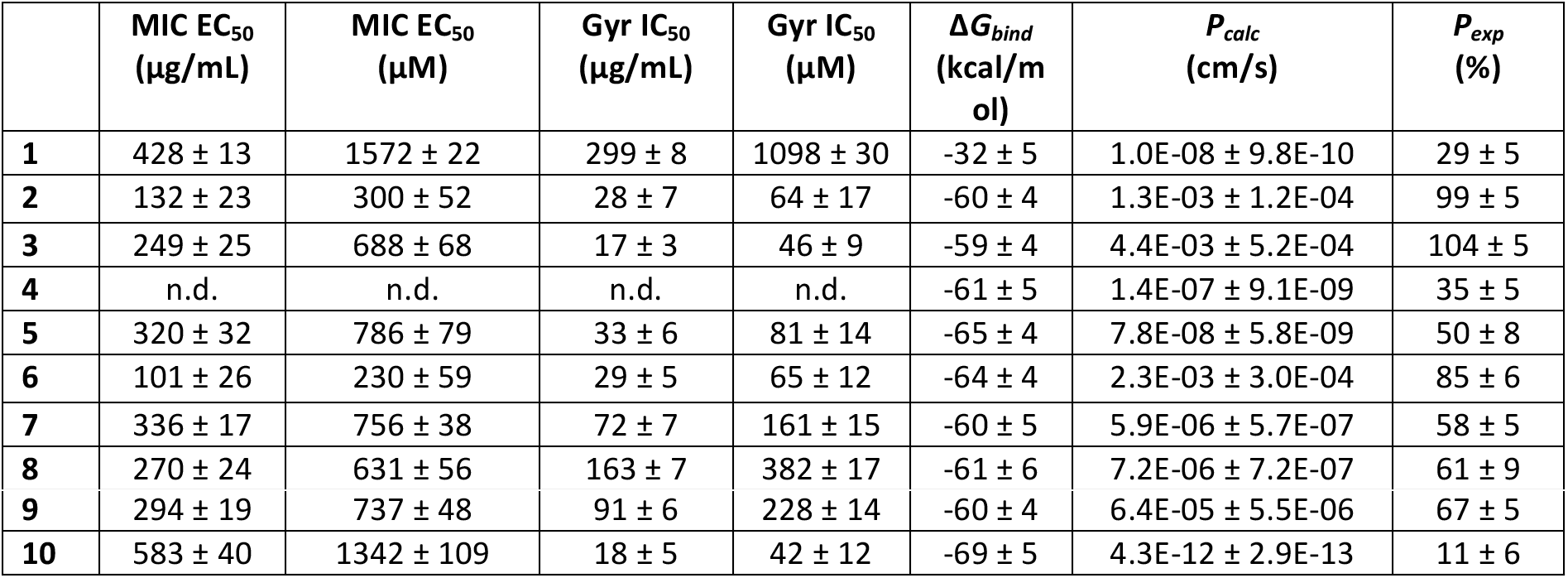
EC_50_ values for growth inhibition (MIC EC_50_), DNA gyrase B supercoiling inhibitory activities (Gyr IC_50_), relative free-energies of binding (Δ*G_bind_*), predicted (*P_calc_*) and experimental (*P_exp_*) OmpF permeabilities for compounds **1-10**.

| | $\text{MIC EC}_{50}$<br>($\mu\text{g/mL}$) | $\text{MIC EC}_{50}$<br>($\mu\text{M}$) | $\text{Gyr IC}_{50}$<br>($\mu\text{g/mL}$) | $\text{Gyr IC}_{50}$<br>($\mu\text{M}$) | $\Delta G_{\text{bind}}$<br>(kcal/mol) | $P_{\text{calc}}$<br>(cm/s) | $P_{\text{exp}}$<br>(%) |
| --- | --- | --- | --- | --- | --- | --- | --- |
| **1** | $428 \pm 13$ | $1572 \pm 22$ | $299 \pm 8$ | $1098 \pm 30$ | $-32 \pm 5$ | $1.0\text{E-}08 \pm 9.8\text{E-}10$ | $29 \pm 5$ |
| **2** | $132 \pm 23$ | $300 \pm 52$ | $28 \pm 7$ | $64 \pm 17$ | $-60 \pm 4$ | $1.3\text{E-}03 \pm 1.2\text{E-}04$ | $99 \pm 5$ |
| **3** | $249 \pm 25$ | $688 \pm 68$ | $17 \pm 3$ | $46 \pm 9$ | $-59 \pm 4$ | $4.4\text{E-}03 \pm 5.2\text{E-}04$ | $104 \pm 5$ |
| **4** | n.d. | n.d. | n.d. | n.d. | $-61 \pm 5$ | $1.4\text{E-}07 \pm 9.1\text{E-}09$ | $35 \pm 5$ |
| **5** | $320 \pm 32$ | $786 \pm 79$ | $33 \pm 6$ | $81 \pm 14$ | $-65 \pm 4$ | $7.8\text{E-}08 \pm 5.8\text{E-}09$ | $50 \pm 8$ |
| **6** | $101 \pm 26$ | $230 \pm 59$ | $29 \pm 5$ | $65 \pm 12$ | $-64 \pm 4$ | $2.3\text{E-}03 \pm 3.0\text{E-}04$ | $85 \pm 6$ |
| **7** | $336 \pm 17$ | $756 \pm 38$ | $72 \pm 7$ | $161 \pm 15$ | $-60 \pm 5$ | $5.9\text{E-}06 \pm 5.7\text{E-}07$ | $58 \pm 5$ |
| **8** | $270 \pm 24$ | $631 \pm 56$ | $163 \pm 7$ | $382 \pm 17$ | $-61 \pm 6$ | $7.2\text{E-}06 \pm 7.2\text{E-}07$ | $61 \pm 9$ |
| **9** | 294 ± 19 | 737 ± 48 | 91 ± 6 | 228 ± 14 | -60 ± 4 | 6.4E-05 ± 5.5E-06 | 67 ± 5 |
| **10** | 583 ± 40 | 1342 ± 109 | 18 ± 5 | 42 ± 12 | -69 ± 5 | 4.3E-12 ± 2.9E-13 | 11 ± 6 |

To rule out the effect of changes to DNA gyrase inhibition, we tested the ability of each compound to inhibit *E. coli* DNA gyrase B in a cell-free context using a DNA supercoiling assay with purified gyrase enzyme. As shown in Fig. 5, measured IC_50_ values for DNA gyrase B inhibition correlate moderately with crude estimates of target binding affinity from MM-PBSA calculations (Pearson *r* = 0.87 however Spearman rho 0.27) but minimally with microbiological EC_50_ values (Pearson *r* = 0.26), suggesting that the improvement in antimicrobial activity was indeed due to enhanced bacterial uptake rather than enhanced biochemical activity once inside the cell. Given the rough approximation involved in MM-PBSA calculations (there exist more accurate methods for estimating experimental binding free energies), these data provide supporting evidence that permeability can be optimized independently of target-binding affinity to improve antimicrobial activity.

**Figure 5.**
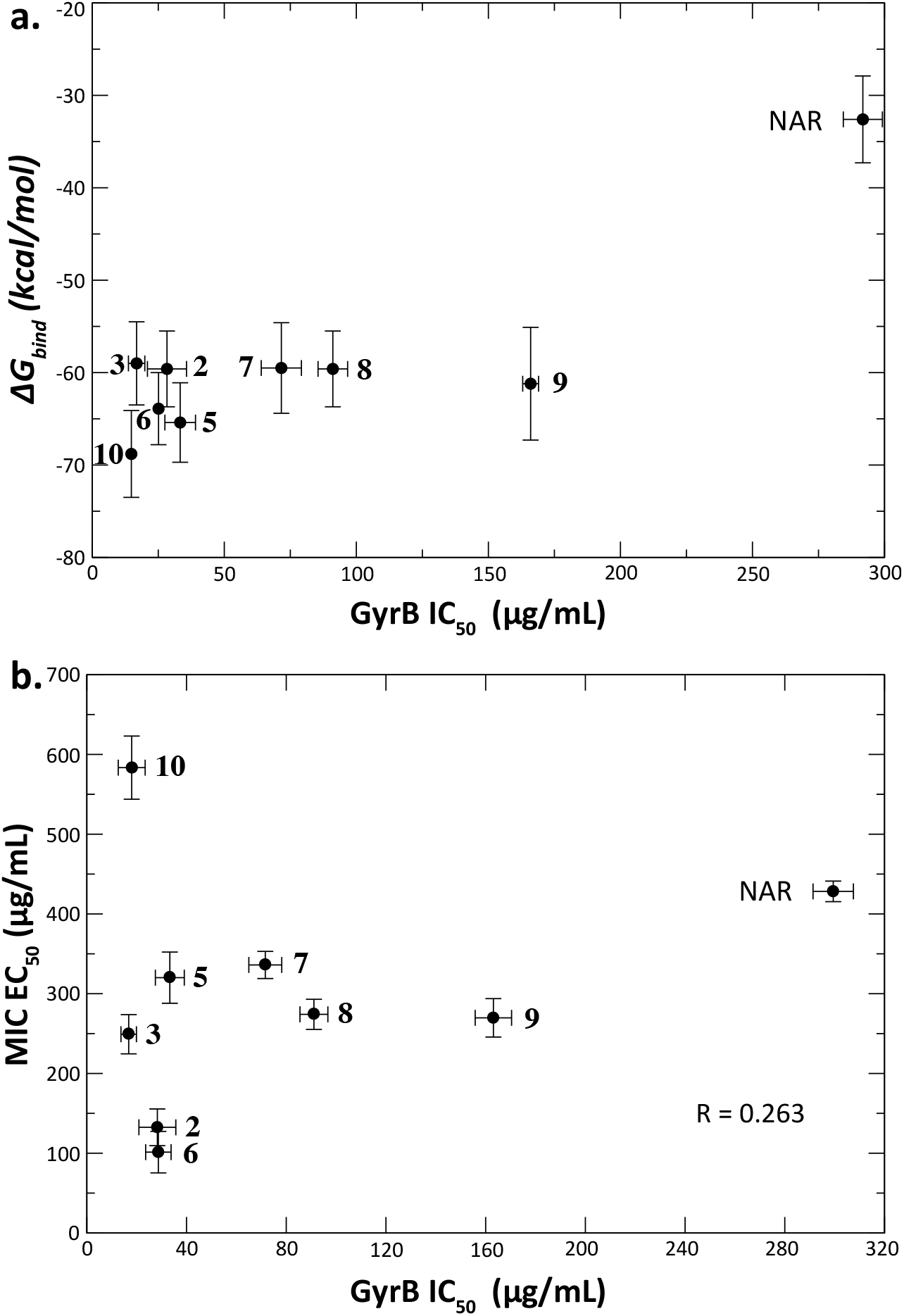
Measured DNA gyrase inhibition correlates moderately well with predicted binding free energy but poorly with antimicrobial activity. Plotted in panel (a) are *E. coli* DNA gyrase B inhibitory concentrations (GyrB IC_50_) and calculated approximate binding free energies. Although these values show a good linear correlation (Pearson r=0.88), the linear fit is dominated by naringinen, and the rank-order correlation is weak (Spearman rho=0.27). Plotted in panel (b) are correlations of DNA gyrase B inhibition with growth inhibitory concentrations (EC_50_) for naringenin (NAR) and all flavanone derivatives except **4**.

## Conclusions

Antimicrobial activity results from a combination of small-molecule uptake, target-binding activity, intracellular degradation, and efflux. Of these, target-binding activity has been the most straightforward to optimize, and small-molecule uptake in Gram-negative bacteria has remained a challenging target. Recent advances in predicting uptake in such bacteria offer the promise to co-optimize candidate molecules for binding and uptake early in the lead generation and optimization process. Machine-learning-based approaches are powerful and computationally efficient in this regard but have thus far been most accurate in well sampled areas of chemical space. Simulation-based prediction offers a more computationally intensive but more flexible approach to optimizing Gram-negative bacterial uptake. Here, we demonstrate that molecular simulations can be used in a prospective manner to select and order derivatives for synthesis and characterization. This simulation-guided engineering was performed for a target and scaffold that the simulation and analysis methodology had not been previously optimized or parameterized for, thus constituting a strong predictive test. This method can be combined with high-throughput prediction of target binding free energies to further enhance computational lead optimization and in the future also coupled to predictions of efflux. We note that compound **D1** (Table S2), with a greatly improved predicted binding ΔG of −80 kcal/mol and moderate permeability of 3.5 × 10^-6^ cm/s, would form a good starting point for such a joint optimization. At least for flavonoid DNA gyrase inhibitors, the molecules used were uptake-dominated in that changes to bacterial membrane permeability capture 86% of the variation in antimicrobial activity. This approach for lead optimization can be readily automated, and we provide code for the requisite analyses. We note that efflux can also be a major determinant to intracellular compound accumulation and antimicrobial activity depending on the compound chemistry. The relative effects of membrane permeability and efflux will vary on the compound set, so our approach should also be evaluated on other compound chemistries to determine generalizability. It nonetheless serves as a proof of concept for increasing antimicrobial activity by optimizing compound uptake.

## Methods

### Compound synthesis

Synthesis of hydrazone derivatives is described in brief below; materials, NMR spectra, yields, and peak assignments are given in the Supplement. 1.5 equiv. of the hydrazine (R-NH-NH_2_) substituent was added to 1 equiv. of naringenin (**1**) dissolved in ethanol containing 0.01 equiv. of 10% acetic acid in ethanol. The mixture was kept stirring and heated until reflux under argon atmosphere until disappearance of the starting compound. The solvent was evaporated, and the residue was extracted with ethyl acetate, dried with anhydrous sodium sulfate, and further purified by preparative thin-layer chromatography (TLC, *n*-hexane:acetone, 1:1) to obtain the corresponding hydrazone (Figures S3-S8). Compound **4** had poor synthetic yield, poor OMV uptake, and high MIC EC_50_ value, so DNA gyrase B inhibition was not determined.

### Docking calculations

The crystallographic structure of DNA Gyrase B ATP binding domain of *Escherichia coli* in complex with a small molecule inhibitor (PDB ID: 4KFG; Figure 1A)^37^, was used in all molecular docking studies. The docking software was selected by evaluating its ability to reproduce the crystallographic pose of the bound inhibitor as follows. Both the Molecular Operating Environment (MOE) v2019.1^38^ and Autodock Vina 1.1.2^39^ packages were able to reproduce the pose of the co-crystallized ligand, although key predicted interactions differed. Both docking protocols reproduced interactions between the methylamino moiety in ring A and Asp73 and also preserving the hydrogen-bond network connecting the pyrimidine ring with Asp73, Arg76 and Thr165 (Figure 1A), but the orientation of the fused octahydropyrrole substituent and its interaction with Asn46 (Figure S2) was better reproduced with MOE than with Vina. Based on these criteria, MOE was chosen for assessment of derivative binding poses.

Sixty-one candidate derivative molecules were built in MOE and minimized using the Amber10:ETH force field. Docking was performed using the Dock module of MOE with default options, using the co-crystallized inhibitor to define the active site. The docking algorithm used was Triangle Matcher with the London dG scoring function (initial docking placement and scoring methodologies, 30 poses) followed by a rigid receptor with a GBVI/WSA dG (post-placement refinement and scoring methodology, 5 poses). The candidate derivatives that maintained the correct orientation inside the ATPase binding pocket and kept key drug-protein interactions observed in the co-crystallized inhibitor were further evaluated for approximate binding free-energies and outer membrane permeabilities.

### Small molecule parameterization

Simulations for binding free-energies were performed using the CHARMM36^40^ and CGenFF force fields^41^, and those for permeability were performed using GROMOS96. Small-molecule topologies for CHARMM36 force field were obtained using the CGenFF server^41^ and converted to GROMACS format using the cgenff_charmm2gmx.py script. For permeability calculations, as in previous work^12^, GROMOS96 parameters for all compounds were assigned using the PRODRG server^42^ with partial charges assigned using the AM1-BCC charge model^43^ in Antechamber 1.27^44^.

### Approximate binding free-energy calculations

Approximate relative free energies of binding were estimated using molecular mechanics-Poisson Boltzmann surface area (MM-PBSA)^45^ calculations based on molecular dynamics (MD) simulations of the ligand-drug complex in GROMACS^46^ using the CHARMM36 and CGenFF force as follows. The 29 compounds passing docking evaluation were simulated via MD starting from their top-ranked binding pose. The system was solvated with TIP3P waters and counter ions was added sufficient to obtain charge neutralization. The system was energy-minimized and then equilibrated for 10 ps under NVT and 1 ns under NPT conditions (Nosé-Hoover thermostat, Parrinello-Rahman barostat), keeping all heavy atoms position-restrained. After equilibration, a 50-ns unrestrained NPT run followed, from which the last 30 ns were considered production. Long-range electrostatics were treated via Particle Mesh Ewald (PME)^47^, with 1.2-nm short-range cut-offs for electrostatic and van der Waals interactions (1.2 nm). SETTLE and LINCS^48^ algorithms were used to constrain all bond lengths. Approximate relative free-energies of binding (total and per residue contributions) were calculated with *g_mmpbsa^49^* for the last 30 ns of the simulation run.

### Prediction of outer-membrane permeability

Predictions of outer-membrane permeability were performed using parameters previously published^12^. Briefly, OmpF was used as a model outer membrane porin and was simulated in a bacterial outer membrane patch consisting of 120 LPS in the outer leaflet and 276 PVPE:68 PVPG:37PVPV-DPG in the inner leaflet. Simulations were performed in GROMACS^46^ using explicit water. Each small molecule compound was first slowly pulled through the porin, and then this pulling simulation was used to seed umbrella sampling and permeability calculations. The single molecule was placed in bulk water at roughly ~1.2 nm from the first OmpF monomer and all overlapping waters removed. The molecule was then pulled to the opposite side of the membrane over 100 ns, using a spring constant of 1000 kJ mol^-1^ nm^-2^ and a pull rate of 0.085 ± 0.005 nm ns^-1^ to obtain an initial path from which starting points for umbrella sampling were chosen at even *z*-intervals. Twenty-six (26) umbrella windows, with a mean width of 0.35 ± 0.02 nm were used to sample the *z* coordinate path and 40 ns of molecular simulation was performed per window (>1 μs total sampling per molecule).

The inhomogeneous solubility-diffusion (ISD) model was used to estimate drug permeabilities as in previous work^12^ using the following equation:

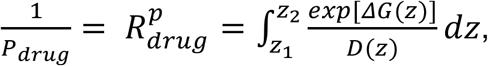

where *P_drug_* is permeability, *R^p^* is resistance to permeation, *β* = 1/*k_B_T*, Δ*G*(*z*) is the potential of mean force from umbrella sampling, *D*(*z*) is the local diffusivity coefficient, also estimated from umbrella sampling, and *z* is the relative position of the molecule along the pore axis. Error estimation for the PMF curves were estimated by bootstrapping over umbrella sampling windows, sampled at intervals equal to the calculated autocorrelation times (*τ_z_*) in each window. PMF curves are plotted in Fig. S9. We also performed validation studies where we rotated compounds 180 degrees at the beginning of the pulling run and used that for umbrella sampling and subsequent PMF and permeability calculation. These validation data yield permeabilities within ~1.1 orders of magnitude, adding an uncertainty term but preserving the log-linear relationship between predicted permeability and measured compound uptake.

### Outer-membrane vesicle swelling assays

Assays were performed as in prior work^12^. Briefly, 5 μL of *E. coli* MG1655, ΔOmpC (kind gift of Linus Sandegren) was inoculated in 5 mL LB broth and incubated overnight at 37 °C/200 rpm. This culture was then seeded into 250 mL of fresh LB broth and incubated for 4 h at 37 °C/100 rpm or to OD_600_ between 1.2 and 1.4. This culture was centrifuged for 20 min at 10,000x *g* and 4 °C. The supernatant was then sterile-filtered through a 0.22 μm filter followed by ultracentrifugation for 4 h at 150,000x *g*, 4 °C. The supernatant was immediately decanted, and the pellet was resuspended in 1 mL MES buffer pH 6.0, for DLS measurements. Size measurements, cryo-electron micrographs, and SDS-PAGE of the OMV preparations are also given in our prior work^12^.

Each compound was dissolved in 100 μL of MES buffer pH 6.0 to a final concentration of 9 mM^12, 50, 51^. To measure vesicle swelling, 6 μL of the OMV suspension was added to 6 μL MES buffer containing 2 μL MES (control) or 2 μL of the desired molecule, to a total volume of 14 μL. The light-scattering autocorrelation function was measured over 60 seconds using a 660 nm laser on an AvidNano W130i instrument (Avid Nano, High Wycombe, UK). All measurements were performed using a quartz cuvette and processed with pUnk 1.0.0.3 (Avid Nano, High Wycombe, UK). At least 3 biological replicates of 4 technical replicates each were performed per compound tested. No unexpected or unusually high safety hazards were encountered.

### DNA gyrase inhibition assays

DNA gyrase supercoiling activity was assessed by measuring the conversion of relaxed plasmid pBR322 DNA to the supercoiled form^52, 53^. Inhibition assays were carried out using an *E. coli* gyrase supercoiling assay kit from Inspiralis (Norwich Research Park, Colney Lane, Norwich. UK) according to manufacturer instructions. Briefly, in 30 μL reaction mixtures containing the DNA gyrase assay buffer [35 mM Tris-HCl (pH 7.5), 24 mM KCl, 4 mM MgCl2, 2 mM DTT, 1.8 mM spermidine, 1 mM ATP, 6.5% (w/v) glycerol and 0.1 mg of bovine serum albumin (BSA)/mL], 150 ng of relaxed pBR322, 2-3 μL of purified DNA gyrase and 3 μL of serial test compounds in a serial 2-fold dilution. 3 μL DMSO was used as control. The mixture was incubated for 30 min at 37 °C, after which the reaction was stopped by adding 30 μL of STEB [40% (w/v) sucrose, 100 mM Tris-HCl (pH 8), 10 mM EDTA, 0.5 mg/mL bromophenol blue] and 30 μL of chloroform/isoamyl alcohol (v:v, 24:1), vortexed briefly and centrifuged for 1 minute. Then, 20 μL of the aqueous upper blue phase was loaded onto a 1 % (w/v) agarose gel in TAE buffer (Tris/EDTA, pH 8.0) and an electrophoresis followed during 2h at 85 V. The gel was then stained with SYBR™ Safe DNA Gel Stain in water (15 min), destained (5-10 min) and visualized under a trans-illuminator. The inhibitory effect of flavonoid on DNA gyrase was assessed by determining the concentration of flavonoid required to inhibit 50% of the supercoiling activity of the enzyme (IC_50_). All determinations were performed as least in triplicate. A total of 1 unit (U) of enzyme activity was defined as the amount of DNA gyrase that converted 150 ng of relaxed pBR322 DNA to the supercoiled form in 30 min.

### Antimicrobial activity

Antibacterial activity was assessed as the concentration that inhibits the growth of 50% of *E. coli* MG1655 strain in broth culture (MIC EC_50_). The EC_50_ of the test compounds was determined by microtiter broth dilution performed in 96-well microplates as described by Wiegand *et al*^54^. Bacterial cultures were prepared by inoculating 5 mL of Mueller Hinton broth with 5 μL of a bacterial glycerol stock followed by incubation at 37 °C/200 rpm for 5 h. The turbidity of the culture was adjusted to A_600_ 0.07 to 0.1 (0.5 McFarland) by dilution with fresh medium. Each compound was tested in twofold dilutions, using ciprofloxacin and novobiocin as internal controls on each plate. 5 μL of bacterial culture was added to 100 μL of compound diluted in Mueller Hinton broth in each well. After incubating the plates for 12-18 h at 37 °C, bacterial growth was examined by measuring the A_600_ optical density (Multiskan FC, Thermo Fisher Scientific). All measurements were performed at least in triplicates. EC_50_ values were also obtained for highly potent and least potent compounds in a ΔTolC E. coli strain (Keio JW5503^55^); comparisons are given in Table S4.

## Acknowledgements

This work was supported by a Coulter Translational Partners award and a Wallenberg Academy Fellowship to P.M.K. The authors thank A. Villamil Giraldo for helpful discussions, M-U. Ferreira for the gift of compounds, D. dos Santos for software, and L. Sandegren for the gift of bacterial strains. Compute time was provided by the HPC2N center supported by the Swedish National Infrastructure for Computing.

