## Supporting data for "Simulation-guided engineering of antibiotics for improved bacterial uptake"

### Table of Contents

**Figure S1.** Predicted binding mode for hydrazones (A), urea (B), guanidine (C) and acetamidine (D) at the *E. coli* gyrase B ATP-binding site.

**Table S1.** Relative free-energies of binding ($\Delta G_{bind}$), contribution per residue, binding mode and predicted permeability coefficients for the different linker moieties and compounds **2**-**4**.

**Table S2.** Relative free-energies of binding ($\Delta G_{bind}$), contribution per residue, binding mode and predicted permeability coefficients for derivatives **D1**-**D29**.

**Figure S2.** Co-crystalized pose of known DNA gyrase B inhibitor and top-ranked docking poses obtained with MOE 2019.01 and Vina 1.1.2.

**Figures S3-S8.** $^{1}$H and $^{13}$C NMR spectra of compounds **5**-**10**.

**Figure S9.** Analysis of calculated permeability profiles. Plotted are diffusion, potentials of mean force (different lines indicate bootstrap-resampling runs), and resistance to permeation profiles from permeability calculations for novobiocin, Trius scaffold and compounds **1**-**10**.

**Table S3.** Physico-chemical parameters for compounds **1**-**10**.

**Table S4**. EC50 ratios for ΔTolC E. coli

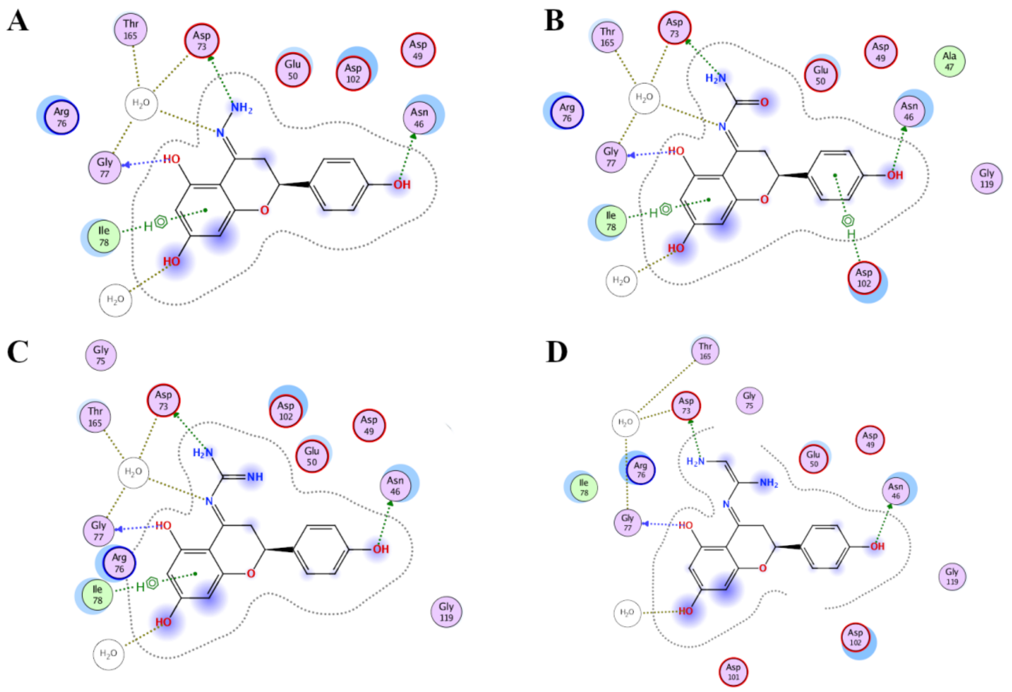

**Figure S1.** Predicted binding mode for hydrazones (A), urea (B), guanidine (C) and acetamidine (D) at the *E. coli* gyrase B ATP-binding site.

**Table S1.** Relative free-energies of binding ($\Delta G_{bind}$), contribution per residue, binding mode and predicted permeability coefficients for the different linker moieties and compounds **2**-**4**.

| Molecule | Scaffold | $\Delta G_{bind}$ | $\Delta G_{bind}$ (per residue) | | | | Bound to D73? | $P_{calc}$ |
|---|---|---|---|---|---|---|---|---|
| | | | N46 | E50 | D73 | R76 | | |
| **naringenin** | =O | -32.6 | -2.0 | -0.5 | -1.4 | -0.9 | Y | $1.00x10^{-08}$ |
| **acetamidine** | =N-C(=CHNH$_2$)-NH$_2$ | -22.5 | -- | -- | 0.1 | -0.4 | N | -- |
| **azine** | =N-N=CH$_2$ | -39.1 | -1.4 | -2.0 | -0.1 | -0.8 | via 5-OH | -- |
| **hydrazone** | =N-NH$_2$ | -44.3 | -2.7 | -3.5 | -2.7 | -0.3 | Y | $1.99x10^{-04}$ |
| **guanidine** | =N-(C=NH)-NH$_2$ | -40.9 | -0.8 | -1.3 | -0.9 | 0.1 | N | -- |
| **urea** | =N-CO-NH$_2$ | -34.9 | -0.8 | -1.7 | -0.6 | 0.2 | N | -- |
| **2** | =N-NH-$C_5H_3BrN$ | -59.6 | -3.8 | -2.4 | -3.3 | -2.3 | Y | $1.30x10^{-03}$ |
| **3** | =N-NH-$C_6H_4$ | -59.0 | -2.7 | -0.2 | -3.0 | -3.5 | Y | $4.37x10^{-03}$ |
| **4** | =N-NH-$C_4H_3N_2$ | -60.8 | -4.7 | -3.3 | -4.5 | -1.7 | Y | $1.36x10^{-07}$ |

$\Delta G_{bind}$, estimated relative free-energy of binding (kcal/mol); $P_{calc}$, predicted outer membrane OmpF porin permeability coefficient (cm/s). Unfavorable residue contributions are depicted in red.

**Table S2.** Relative free-energies of binding ($\Delta G_{bind}$), contribution per residue, binding mode and predicted permeability coefficients for derivatives **D1**-**D29.**

| Mol | Scaffold | $\Delta G_{bind}$ | $\Delta G_{bind}$ (per residue) | | | | Bound to D73? | $P_{calc}$ |
|---|---|---|---|---|---|---|---|---|
| | | | N46 | E50 | D73 | R76 | | |
| **D1** | $=N-NH-(3-SO_2NH_2)-C_6H_4$ | -80.5 | -3.5 | -- | -5.3 | -4.5 | Y | $3.53x10^{-06}$ |
| **D2 (5)** | $=N-NH-(3-NO_2)-C_6H_4$ | -65.4 | -2.7 | -0.9 | -4.2 | -3.4 | Y | $7.80x10^{-08}$ |
| **D3** | $=N-NH-(3-CO_2)-C_6H_4$ | -22.4 | -0.1 | 0.3 | 0.3 | -0.9 | N | -- |
| **D4** | $=N-NH-(2,5-diCl)-C_6H_3$ | -64.8 | -3.1 | -1.2 | -4.3 | -3.0 | Y | $1.14x10^{-07}$ |
| **D5 (10)** | $=N-NH-(3-CO_2CH_2CH_3)-C_6H_4$ | -68.8 | -3.5 | -0.9 | -3.2 | -3.3 | Y | $4.31x10^{-12}$ |
| **D6** | $=N-NH-CH_2-(3-CH_3)-C_6H_4$ | -43.1 | -1.6 | -3.3 | -0.2 | -0.6 | N | -- |
| **D7** | $=N-NH-C_9H_7N$ | -19.7 | 0.1 | 10.5 | 4.1 | -1.3 | N | -- |
| **D8** | $=N-NH-(3-CN)-C_6H_4$ | -69.7 | -3.3 | -0.9 | -4.0 | -3.2 | Y | $1.40x10^{-06}$ |
| **D9** | $=N-NH-CH_2-C_6H_4$ | -38.3 | -1.2 | -0.6 | -0.4 | 0.4 | N | -- |
| **D10** | $=N-NH-CH_2-(2-CH_3)-C_6H_4$ | -49.0 | -3.2 | 1.2 | 1.1 | -1.2 | Y | $9.76x10^{-05}$ |
| **D11 (7)** | $=N-NH-CH_2-(3,4-diCl)-C_6H_3$ | -59.5 | -2.2 | 1.4 | -4.5 | -2.6 | Y | $5.86x10^{-06}$ |
| **D12** | $=N-NH-CH_2-(3-Cl)-C_6H_4$ | -48.6 | -1.8 | -2.4 | -- | -0.9 | N | -- |
| **D13** | $=N-NH-(2-NO_2)-C_6H_4$ | -62.9 | -2.5 | 0.8 | 1.2 | -1.2 | N | -- |
| **D14** | $=N-NH-CH_2-(4-F)-C_6H_4$ | -29.6 | -1.3 | 1.2 | -- | -3.1 | N | -- |
| **D15** | $=N-NH-C_9H_8$ | -58.2 | -3.0 | 1.1 | 2.0 | -1.3 | Y | $7.05x10^{-09}$ |
| **D16** | $=N-NH-(2-NH_2, 3-SH)-C_2N_3$ | -65.4 | -2.1 | -1.1 | -5.4 | -1.1 | Y | $3.60x10^{-06}$ |
| **D17** | $=N-NH-(2-OH)-C_5H_8$ | -54.6 | -3.0 | -0.1 | 0.6 | -1.3 | via $2H_2O$ | $4.45x10^{-08}$ |
| **D18 (6)** | $=N-NH-(3-Br)-C_6H_4$ | -63.9 | -3.6 | -1.5 | -3.5 | -2.9 | Y | $2.29x10^{-03}$ |
| **D19** | $=N-NH-(4-CO_2)-C_6H_4$ | -40.1 | -7.9 | 1.6 | 2.5 | -5.2 | Y | $1.57x10^{-09}$ |
| **D20 (8)** | $=N-NH-(4-CH_3)-C_9H_6N$ | -61.2 | -3.5 | -0.5 | 0.7 | -0.3 | Y | $7.21x10^{-06}$ |
| **D21** | $=N-NH-C_3H_3N_2$ | -62.0 | -1.8 | 0.9 | -1.2 | -1.3 | Y | $9.03x10^{-16}$ |
| **D22** | $=N-NH-(2-CH_3)-C_3H_3N_2$ | -62.9 | -3.7 | -1.1 | -3.4 | -3.3 | Y | $1.27x10^{-05}$ |
| **D23** | $=N-NH-(2-CH_2CH_3)-C_3H_3N_2$ | -68.1 | -4.3 | -0.9 | -3.9 | -3.5 | Y | $8.75x10^{-06}$ |
| **D24** | $=N-NH-(2-CH_3-5-CF_3)-C_3H_2N_2$ | -61.9 | -3.1 | -1.2 | -3.8 | -3.0 | Y | $5.13x10^{-06}$ |
| **D25** | $=N-NH-(2,4-diCH_3-4-NO_2)-C_3H_2N_2$ | -52.0 | -2.0 | -0.2 | 0.8 | -0.2 | via 5-OH | $7.09x10^{-05}$ |
| **D26** | $=N-NH-(4-Br)-C_3H_3N_2$ | -64.6 | -4.0 | -5.4 | -4.6 | -1.0 | Y | $2.26x10^{-07}$ |
| **D27** | $=N-NH-(3-Cl)-C_3H_2N_2$ | -64.1 | -5.7 | -9.1 | -11.8 | -0.2 | Y | $4.81x10^{-08}$ |
| **D28 (9)** | $=N-NH-(3-Cl)-C_4H_2N_2$ | -59.6 | -5.3 | -4.3 | -4.6 | -1.2 | Y | $6.42x10^{-05}$ |
| **D29** | $=N-NH-C_8H_5N_2$ | -48.5 | -0.7 | -7.6 | 1.1 | -0.9 | Y | $1.02x10^{-09}$ |

$\Delta G_{bind}$, estimated relative free-energy of binding (kcal/mol); $P_{calc}$, predicted outer membrane OmpF porin permeability coefficient. Unfavorable residue contributions are depicted in red. Compounds highlighted in green were synthesized and experimentally evaluated.

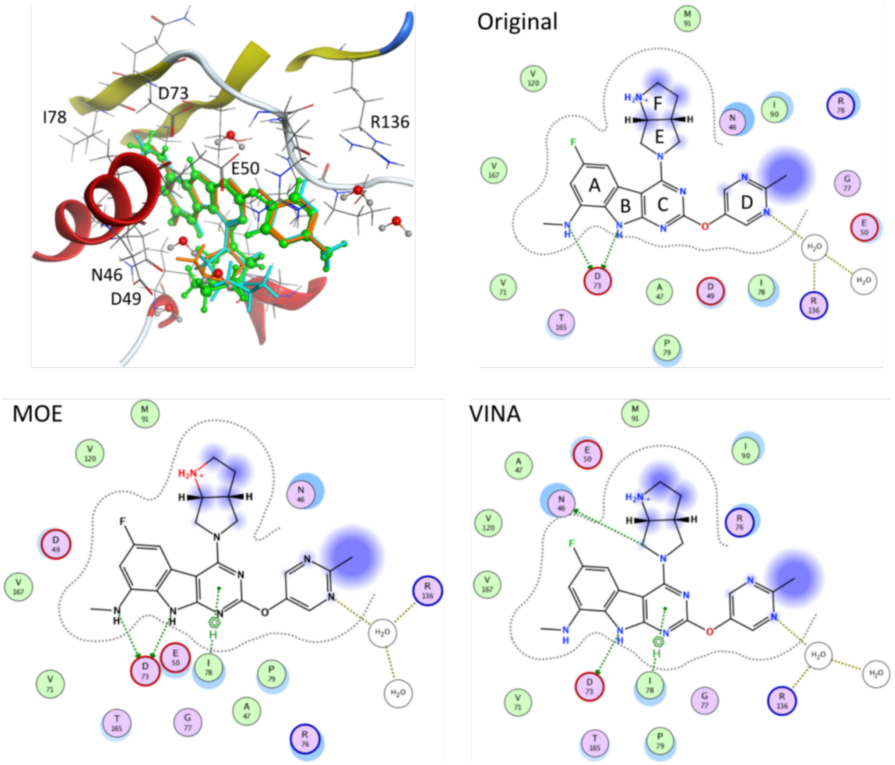

**Figure S2.** Co-crystalized pose of known DNA gyrase B inhibitor and top-ranked docking poses obtained with MOE 2019.01 and Vina 1.1.2.

(*E*)-2-(4-hydroxyphenyl)-4-(2-(3-nitrophenyl)hydrazineylidene)chromane-5,7-diol (**5**), amorphous, light orange powder, from the reaction of naringenin (**1**) with 3-nitrophenylhydrazine (30.9 mg; 51.6% yield); $^{1}$H NMR (500 MHz, Acetone-$D_6$) $\delta$ 7.76 (1H, t, $J$ = 2.3 Hz, H-2”), 7.61 (1H, dd, $J$ = 8.1, 1.5 Hz, H-4”), 7.53 (1H, t, $J$ = 8.1 Hz, H-5”), 7.35 (3H, m, H-2’, H-6”), 6.82 (2H, d, $J$ = 8.5 Hz, H-3’), 5.97 (1H, d, $J$ = 2.3 Hz, H-6), 5.89 (1H, d, $J$ = 2.3 Hz, H-8), 5.11 (1H, dd, $J$ = 12.1, 3.1 Hz, H-2), 3.29 (1H, dd, $J$ = 16.9, 3.1 Hz, H-3β), 2.88 (1H, dd, $J$ = 16.9, 12.1 Hz, H-3α) ppm; $^{13}$C NMR (125 MHz, Acetone-$D_6$) $\delta$ 160.3 (C-7), 159.3 (C-9), 158.2 (C-5), 157.6 (C-4’), 148.8 (C-3”), 147.2 (C-4), 146.1 (C-1”), 130.7 (C-1’, C-5”), 128.1 (C-2’), 118.0 (C-6”), 115.2 (C-2”), 113.2 (C-4”), 105.8 (C-2”), 98.5 (C-10), 96.6 (C-8), 95.3 (C-6), 75.9 (C-2), 32.0 (C-3) ppm.

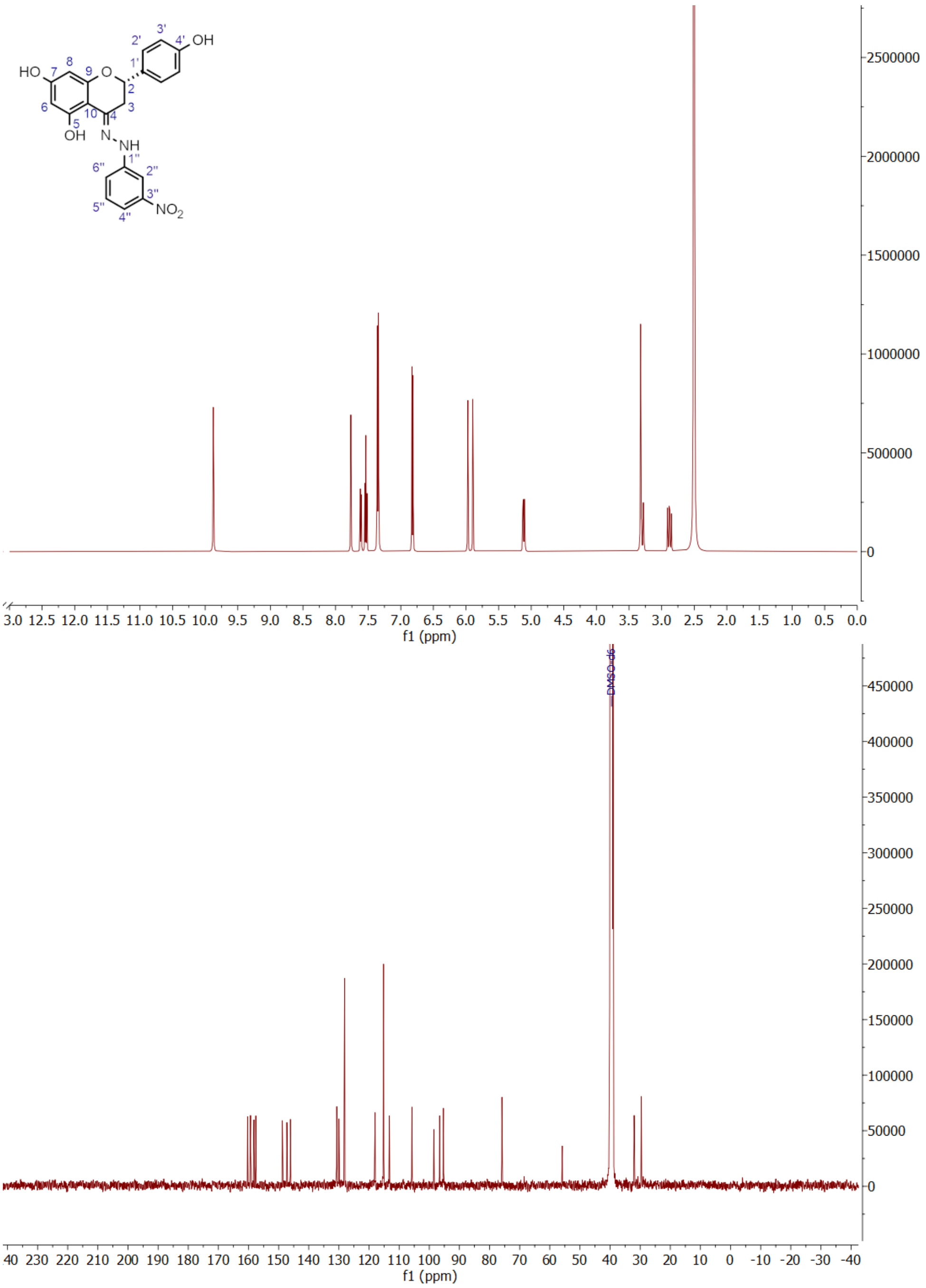

**Figure S3.** $^{1}$H and $^{13}$C NMR spectra of compound **5**.

(*E*)-4-(2-(3-bromophenyl)hydrazineylidene)-2-(4-hydroxyphenyl)chromane-5,7-diol (**6**), amorphous, brown powder, from the reaction of naringenin (**1**) with 3-bromophenyl hydrazine (7.2 mg; 11% yield);

$^{1}$H NMR (500 MHz, Acetone-$D_6$) $\delta$ 7.40 (2H, d, *J* = 8.5 Hz, H-2’), 7.23 (1H, d, *J* = 2.2 Hz, H-2”), 7.20 (1H, t, *J* = 8.0 Hz, H-5’’), 7.03 (1H, dd, *J* = 8.0, 2.0 Hz, H-4’’), 6.98 (1H, dd, *J* = 8.0, 2.0 Hz, H-6’’), 6.91 (2H, d, *J* = 8.5 Hz, H-3’), 6.06 (1H, d, *J* = 2.4 Hz, H-8), 5,98 (1H, d, *J* = 2.4 Hz, H-6), 5.11 (1H, dd, *J* = 12.1, 3.1 Hz, H-2), 3.28 (1H, dd, *J* = 16.8, 3.1 Hz, H-3β), 2.98 (1H, dd, *J* = 16.8, 12.1 Hz, H-3α) ppm; $^{13}$C NMR (125 MHz, Acetone-$D_6$) $\delta$ 161.1 (C-7), 161.0 (C-9), 159.4 (C-5, C-4’), 147.6 (C-4, C-4”), 131.6 (C-1’, C-5”), 123.6 (C-3”), 128.7 (C-2’), 123.0 (C-6’’), 116.1 (C-3’), 115.6 (C-2’’), 112.1 (C-4’’), 99.9 (C-10), 97.7 (C-8), 96.1 (C-6), 77.3 (C-2), 32.8 (C-3) ppm.

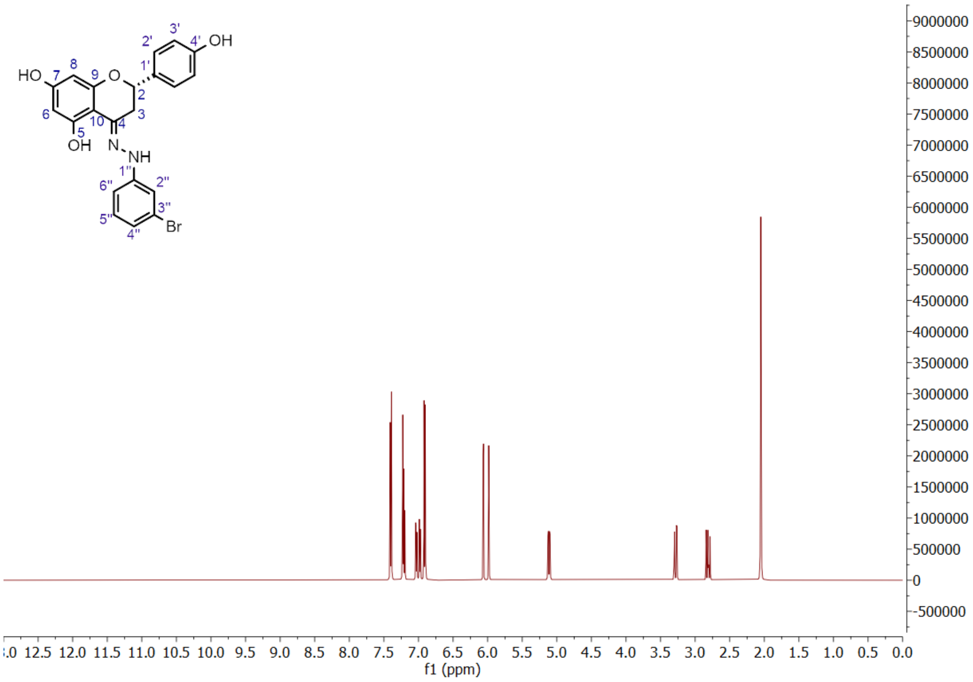

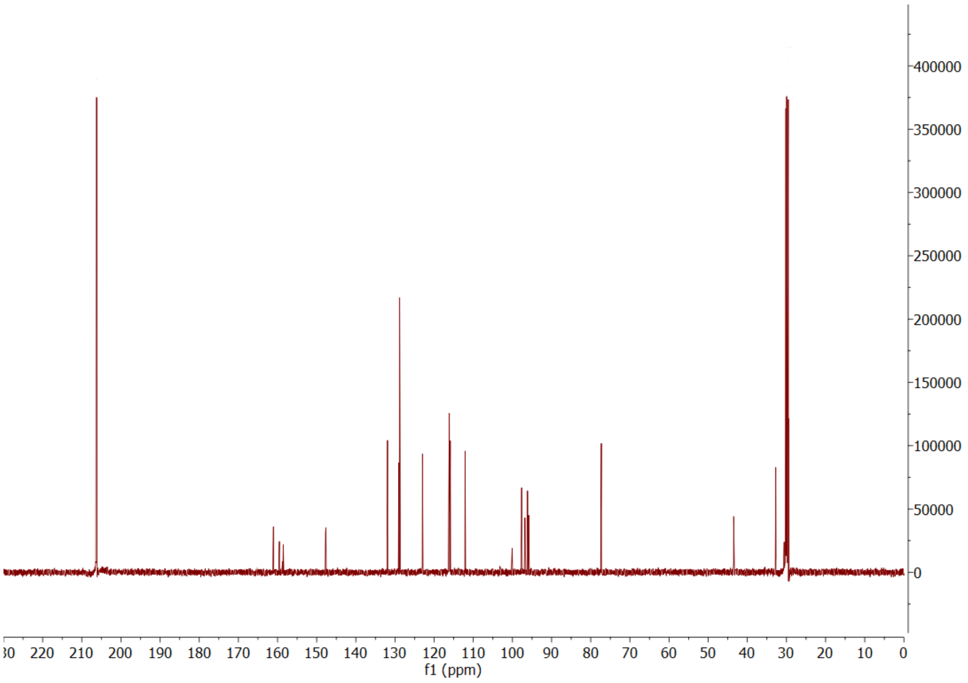

**Figure S4.** $^{1}$H and $^{13}$C NMR spectra of compound **6**.

(*E*)-4-(2-(3,4-dichlorobenzyl)hydrazineylidene)-2-(4-hydroxyphenyl)chromane-5,7-diol (**7**), amorphous, yellow powder, from the reaction of naringenin (**1**) with 3,4-dichlorobenzyl hydrazine (18.9 mg; 30%

yield); [1]H NMR (500 MHz, Acetone-$D_6$) $\delta$ 7.62 (1H, d, *J* = 2.0 Hz, H-3”), 7.53 (1H, d, *J* = 8.2 Hz, H-6”), 7.35 (2H, d, *J* = 8.7 Hz, H-2’), 6.90 (1H, d, *J* = 8.2 Hz, H-7”), 6.88 (1H, d, *J* = 8.7 Hz, H-3’), 5.93 (1H, d, *J* = 2.4 Hz, H-6), 5.90 (1H, d, *J* = 2.4 Hz, H-8), 5.01 (1H, dd, *J* = 11.8, 3.2 Hz, H-2), 4.37 (1H, d, *J* = 4.2 Hz,p H-1”), 3.17 (1H, dd, *J* = 16.8, 3.2 Hz, H-3β), 2.70 (1H, dd, *J* = 16.8, 11.8 Hz, H-3α) ppm; [13]C NMR (125 MHz, Acetone-$D_6$) $\delta$ 167.6 (C-7), 165.3 (C-9), 161.5 (C-5), 159.2 (C-4’), 148.4 (C-4), 142.0 (C-5”), 132.0 (C-7”), 131.2 (C-1’), 130.8 (C-3”), 129.2 (C-4”), 129.3 (C-6”), 128.8 (C-2’), 128.5 (C-2”), 116.4 (C-3’), 100.2 (C-10), 97.4 (C-8), 95.7 (C-6), 77.6 (C-2), 53.8 (C-1”), 32.2 (C-3) ppm.

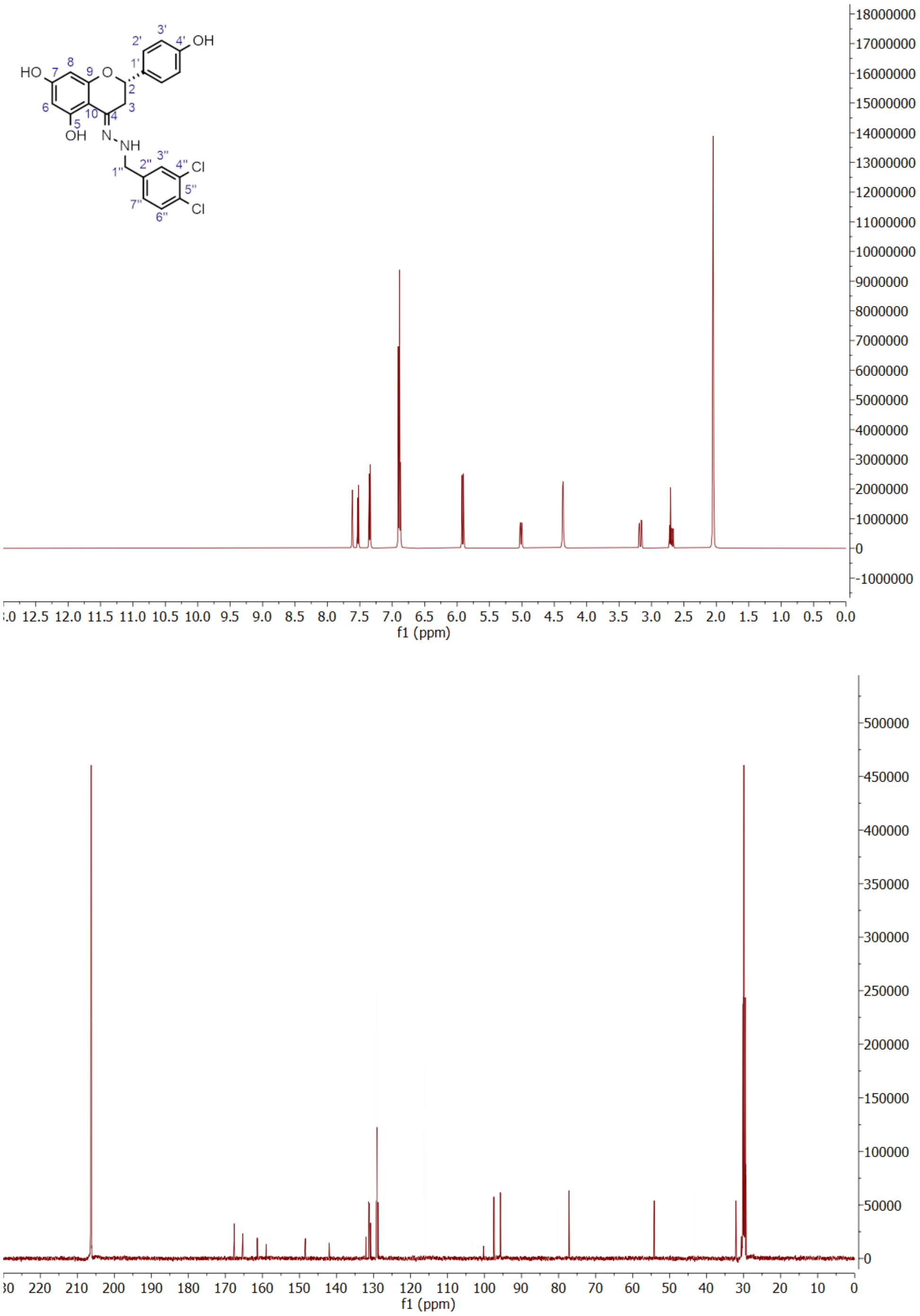

**Figure S5.** [1]H and [13]C NMR spectra of compound **7**.

(*E*)-2-(4-hydroxyphenyl)-4-(2-(4-methylquinolin-2-yl)hydrazineylidene)chromane-5,7-diol (**8**), amorphous, brown powder, from the reaction of naringenin (**1**) with 4-methylquinolin-2-hydrazine

(15.2 mg; 24% yield); $^{1}H$ NMR (500 MHz, Acetone-$D_6$) $\delta$ 7.99 (1H, d, $J$ = 7.8 Hz, H-7”), 7.79 (1H, d, $J$ = 7.8 Hz, H-4”), 7.74 (1H, t, $J$ = 7.8 Hz, H-6”), 7.63 (1H, t, $J$ = 7.8 Hz, H-5”), 7.57 (1H, s, H-2”), 7.40 (2H, d, $J$ = 8.2 Hz, H-2’), 6.90 (2H, d, $J$ = 8.2 Hz, H-3’), 5.96 (1H, d, $J$ = 2.1 Hz, H-6), 5.95 (1H, d, $J$ = 2.1 Hz, H-8), 5.46 (1H, dd, $J$ = 12.8, 3.0 Hz, H-2), 3.18 (1H, dd, $J$ = 17.3, 3.0 Hz, H-3β), 2.82 (3H, s, Me-10”), 2.73 (1H, dd, $J$ = 17.3, 12.8 Hz, H-3α) ppm; $^{13}C$ NMR (125 MHz, Acetone-$D_6$) $\delta$ 167.6 (C-7), 164.8 (C-9), 158.7 (C-4’), 150.8 (C-5), 148.1 (C-8”), 146.9 (C-1”), 145.6 (C-4), 142.0 (C-3”), 131.0 (C-1’), 129.1 (C-2’), 128.5 (C-4”), 128.4 (C-6”), 127.4 (C-5”), 123.8 (C-7”), 122.8 (C-9”), 117.3 (C-2”), 116.1 (C-3’), 97.6 (C-10), 96.8 (C-8), 95.8 (C-6), 79.3 (C-2), 42.6 (C-3), 18.5 (C-10”) ppm.

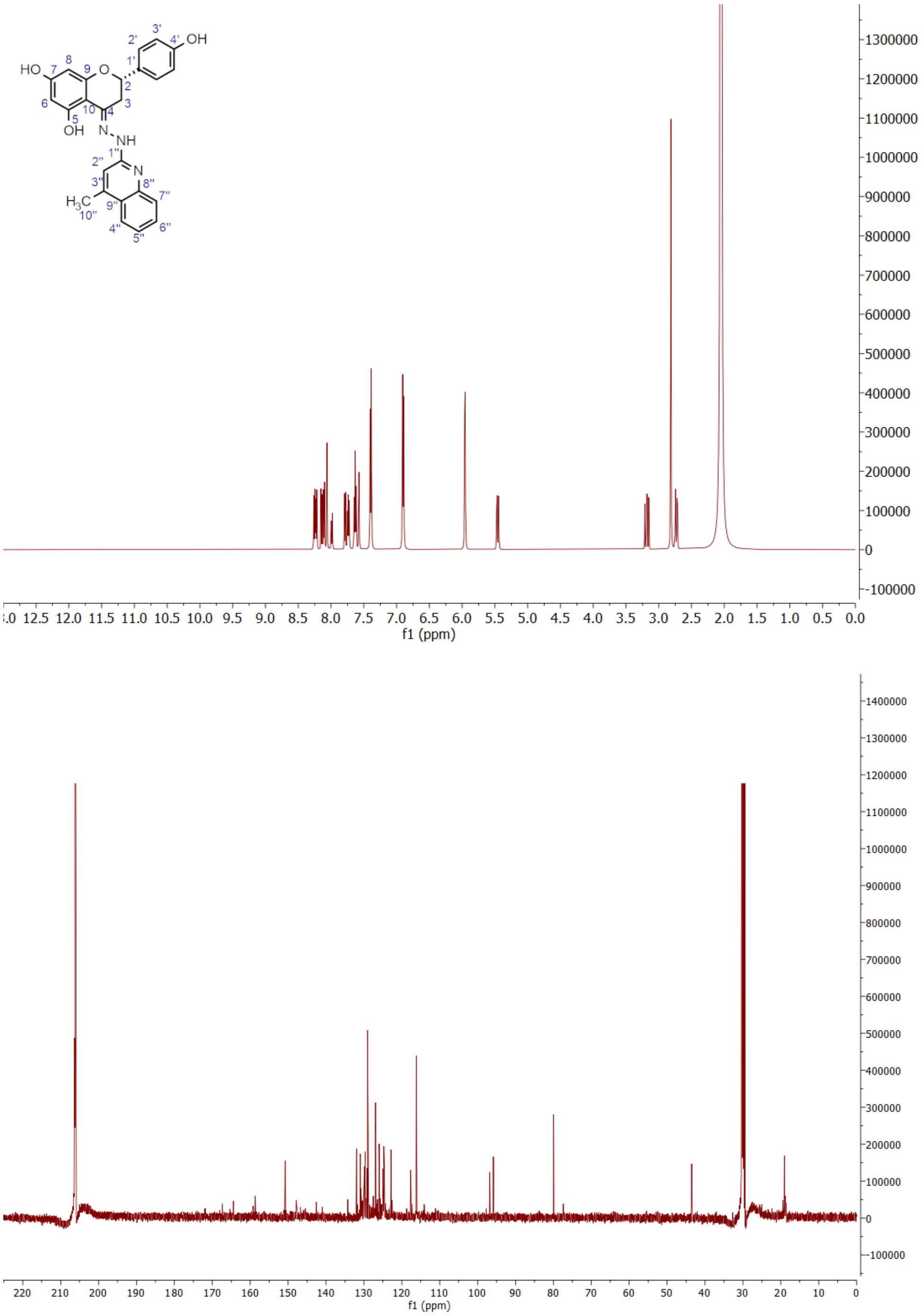

**Figure S6.** $^{1}H$ and $^{13}C$ NMR spectra of compound **8**.

(*E*)-4-(2-(6-chloropyridazin-3-yl)hydrazineylidene)-2-(4-hydroxyphenyl)chromane-5,7-diol (**9**), amorphous, brown powder, from the reaction of naringenin (**1**) with 3-chloro-6-hydrazinyl pyridazine (3.7 mg; 6% yield); $^{1}H$ NMR (500 MHz, Acetone-$D_6$) $\delta$ 7.58 (1H, d, $J$ = 9,2 Hz, H-3”), 7.42 (2H, d, $J$ = 8.0

Hz, H-2’), 7.35 (1H, d, $J$ = 9.2 Hz, H-2”), 6.91 (2H, d, $J$ = 8.0 Hz, H-3’), 6.06 (1H, d, $J$ = 2.3 Hz, H-8), 6.00 (1H, d, $J$ = 2.3 Hz, H-6), 5.16 (1H, br d, $J$ = 11.7 Hz, H-2), 3.46 (1H, br d, $J$ = 16.6 Hz, H-3β), 2.98 (1H, dd, $J$ = 16.6, 11.7 Hz, H-3α) ppm; $^{13}$C NMR (125 MHz, Acetone-D$_6$) $\delta$ 161.6 (C-7), 161.3 (C-9), 159.6 (C-5), 158.8 (C-4’), 158.4 (C-4”), 131.6 (C-1”), 130.4 (C-3”), 128.8 (C-2’, C-2”), 116.1 (C-3’), 99.9 (C-10), 97.7 (C-8), 96.1 (C-6), 77.3 (C-2), 34.9 (C-3) ppm.

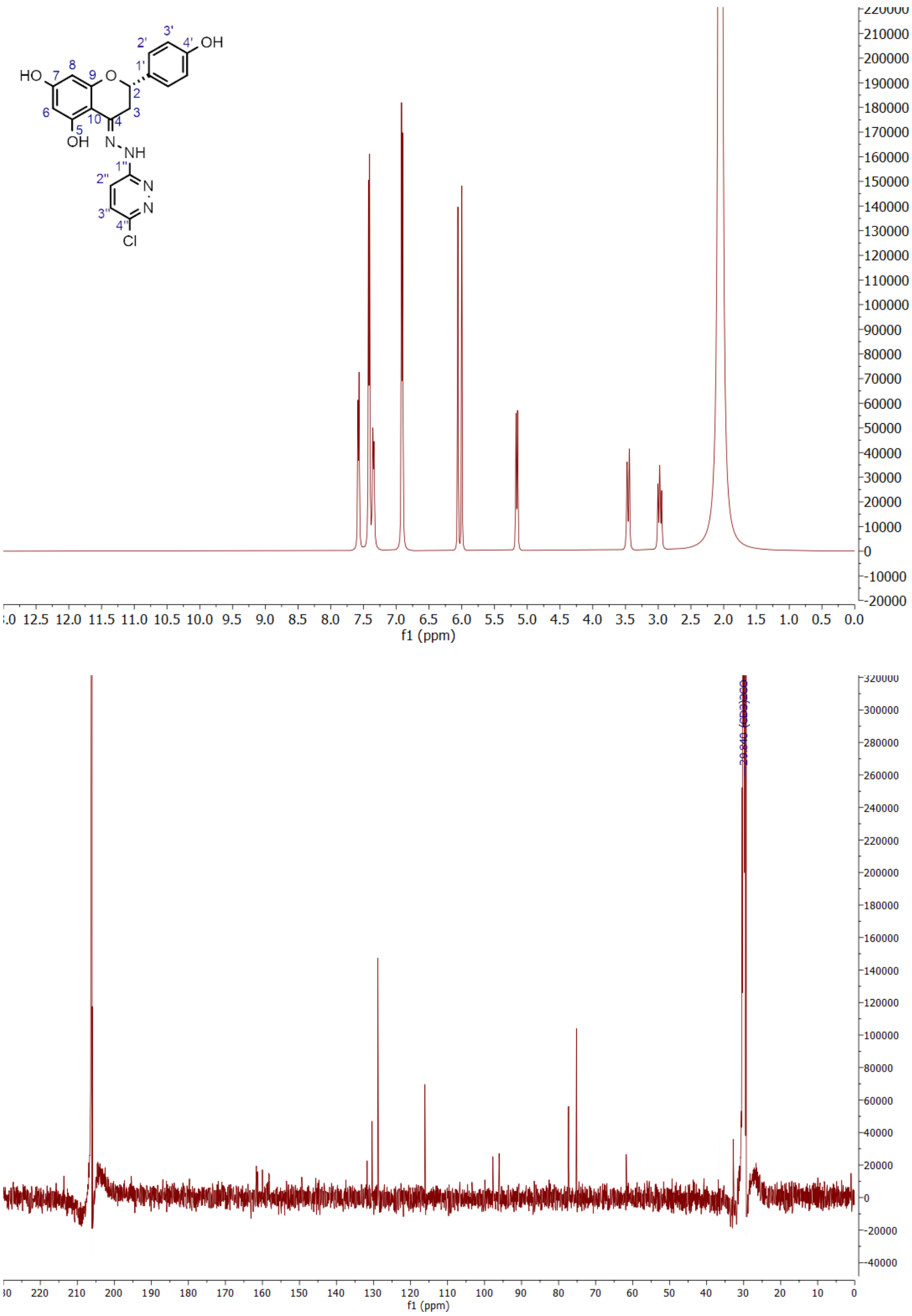

**Figure S7.** $^{1}$H and $^{13}$C NMR spectra of compound **9**.

Ethyl (*E*)-3-(2-(5,7-dihydroxy-2-(4-hydroxyphenyl)chroman-4-ylidene)hydrazineyl) benzoate (**10**), amorphous, brown powder from the reaction of naringenin (**1**) with ethyl 3-hydrazinylbenzoate (8.4 mg; 13% yield); $^{1}$H NMR (500 MHz, Acetone-D$_6$) $\delta$ 7.74 (1H, d, $J$ = 2.1 Hz, H-2”), 7.49 (1H, d, $J$ = 7.9 Hz,

H-4”), 7.41 (2H, d, *J* = 8.4 Hz, H-2’), 7.40 (1H, t, *J* = 7.9 Hz, H-5”), 7.29 (1H, dd, *J* = 7.9, 2.3 Hz, H-6”), 6.91(2H, d, *J* = 8.4 Hz, H-3’), 6.07 (1H, d, *J* = 2.4 Hz, H-8), 5.99 (1H, d, *J* = 2.4 Hz, H-6), 5.12 (1H, dd, *J* = 12.0, 3.0 Hz, H-2), 4.33 (2H, q, *J* = 7.1 Hz, H-8”), 3.35 (1H, dd, *J* = 16.7, 3.0 Hz, H-3β), 2.85 (1H, dd, *J* = 16.7, 12.0 Hz, H-3α), 1.36 (3H, t, *J* = 7.1 Hz, H-9”) ppm; $^{13}$C NMR (125 MHz, Acetone-$D_6$) $\delta$ 167.4 (C-7”), 161.6 (C-7), 161.0 (C-9), 159.5 (C-5), 158.5 (C-4’), 147.2 (C-4), 146.7 (C-1”), 146.5 (C-5”), 131.9 (C-1’), 129.0 (C-5”), 128.9 (C-2’), 121.2 (C-4”), 117.2 (C-6”), 115.9 (C-3’), 113.9 (C-2”), 98.0 (C-10), 97.2 (C-8), 95.9 (C-6), 77.2 (C-2), 61.3 (C-8”), 32.0 (C-3), 14.4 (C-9”) ppm.

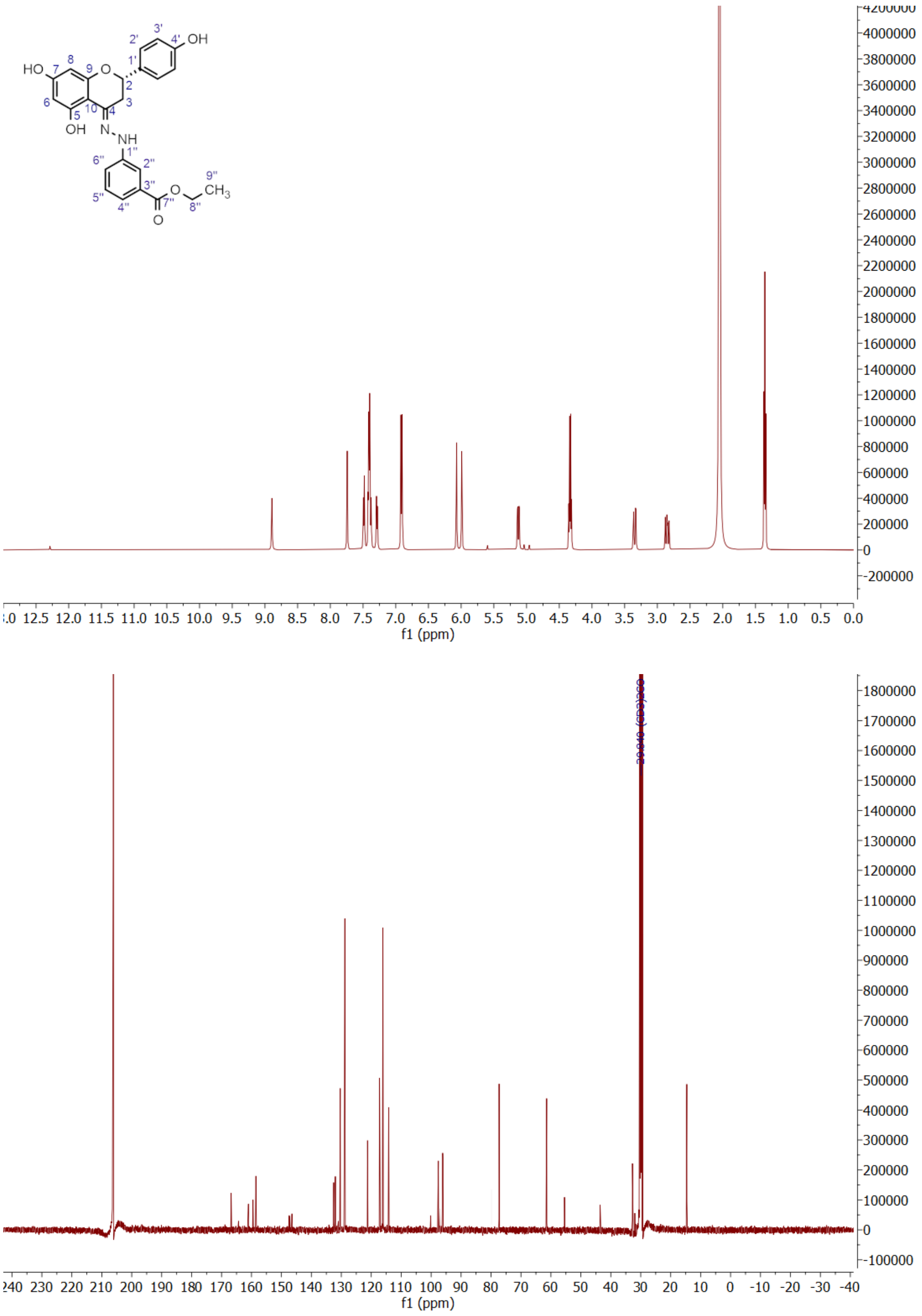

**Figure S8.** $^{1}$H and $^{13}$C NMR spectra of compound **10**.

**Novobiocin**

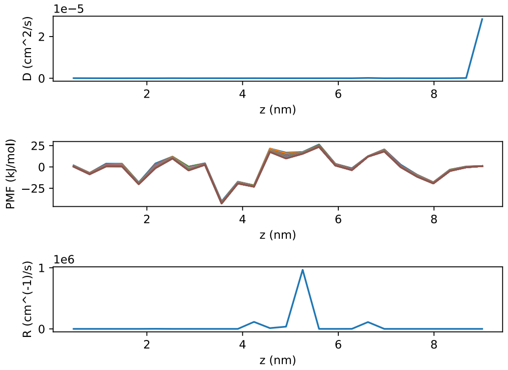

**Trius scaffold**

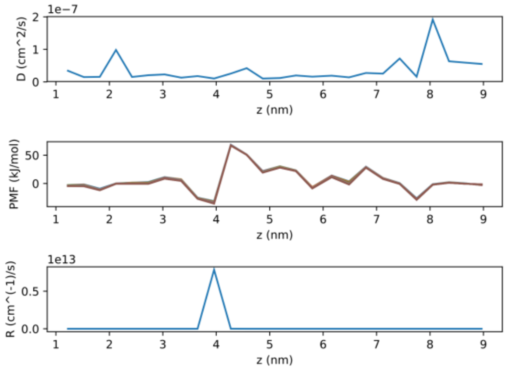

**1**

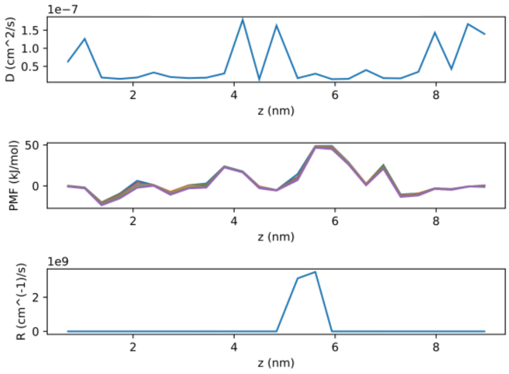

**2**

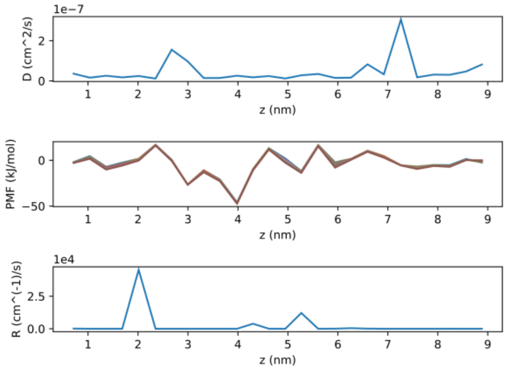

**3**

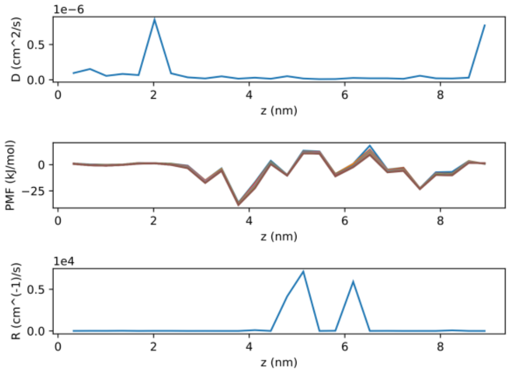

**4**

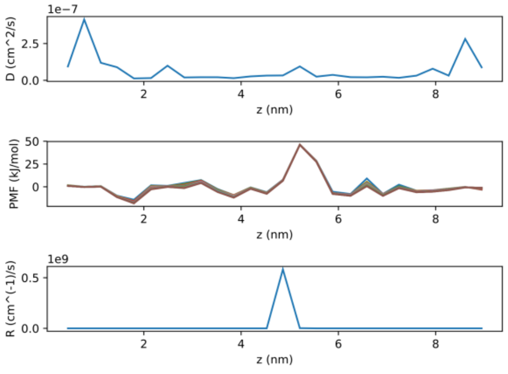

**5**

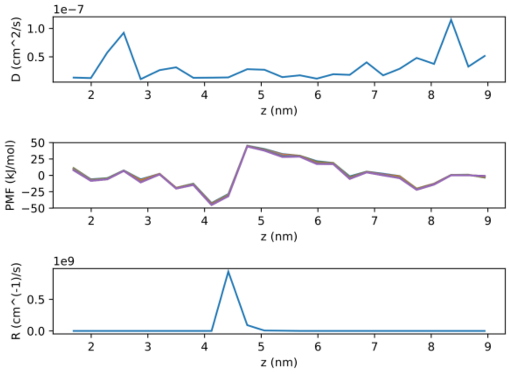

**6**

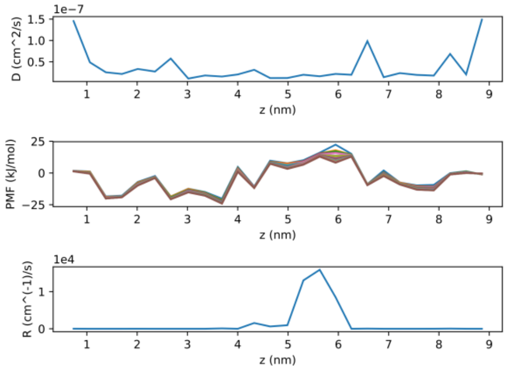

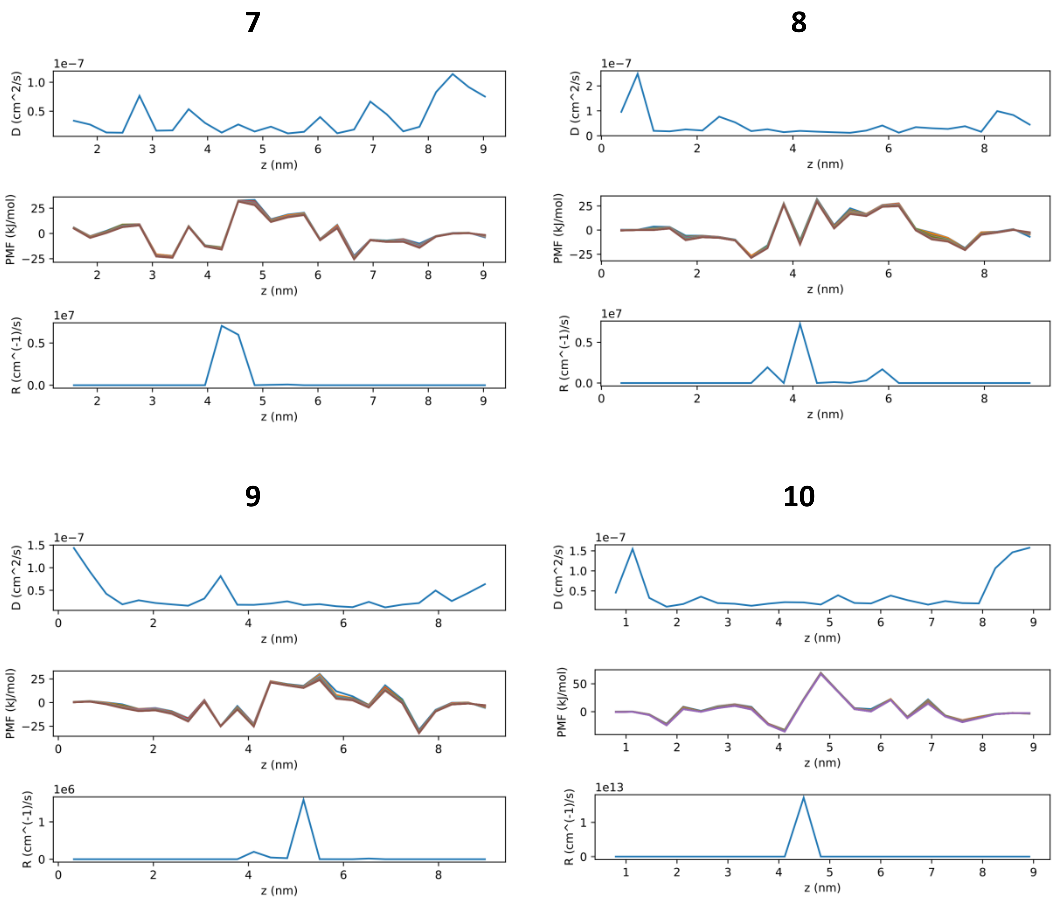

**Figure S9.** Analysis of calculated permeability profiles. Plotted are diffusion (*D*), potentials of mean force (PMF, different lines indicate bootstrap-resampling runs), and resistance to permeation (*R*) profiles from permeability calculations for novobiocin, Trius scaffold and compounds **1-10.**

**Table S3.** Physico-chemical parameters for compounds **1**-**10**.

| | **MW** | **HB don/acc** | **logS** | **log*P*** | **pI** | **dipole (Debye)** | **mPA (Å²)** | **MPA (Å²)** | **vdW_v (Å³)** | **PSA (Å²)** |
|---|---|---|---|---|---|---|---|---|---|---|
| **1** | 272 | 3 / 5 | -2.67 (high) | 2.84 | -- | 4.92 | 4.43 | 7.11 | 229.1 | 87.0 |
| **2** | 442 | 4 / 7 | -4.49 (moderate) | 4.45 | 5.73 | 5.99 | 6.39 | 8.87 | 328.0 | 107.2 |
| **3** | 362 | 4 / 6 | -4.22 (moderate) | 4.30 | 6.00 | 3.02 | 7.27 | 8.08 | 314.2 | 94.3 |
| **4** | 364 | 4 / 8 | -2.43 (high) | 2.46 | 5.56 | 6.27 | 7.01 | 8.11 | 305.8 | 120.1 |
| **5** | 407 | 4 / 8 | -4.83 (low) | 4.24 | 5.58 | 10.77 | 7.22 | 8.92 | 337.2 | 137.5 |
| **6** | 441 | 4 / 6 | -5.11 (low) | 5.07 | 5.79 | 3.59 | 7.28 | 8.34 | 332.4 | 94.3 |
| **7** | 445 | 4 / 6 | -5.39 (low) | 5.06 | 5.12 | 3.80 | 6.63 | 8.86 | 358.9 | 94.3 |
| **8** | 427 | 4 / 7 | -5.72 (low) | 5.57 | 6.60 | 5.48 | 6.69 | 7.71 | 369.9 | 107.2 |
| **9** | 399 | 4 / 8 | -4.85 (low) | 3.52 | 5.43 | 8.38 | 6.53 | 8.69 | 319.4 | 120.1 |
| **10** | 434 | 4 / 7 | -4.67 (low) | 4.66 | 5.92 | 8.78 | 6.98 | 9.23 | 379.1 | 120.6 |

MW, molecular weight; HB don/acc, number of hydrogen-bond donors and acceptors; logS, aqueous solubility; log*P*, octanol-water partition coefficient; pI, isoelectric point; mPA, minimum projection area; MPA, maximum projection area; PSA, polar surface area. All parameters were calculated with MarvinSketch v19.16.

**Table S4**. $EC_{50}$ ratios for ΔTolC E. coli. Lower ratios mean that inhibition is more potent without TolC present and thus the compound is a better TolC substrate.

| **Molecule** | **Ratio ΔTolC/ΔOmpC** |
|---|---|
| **2** | 0.02 |
| **3** | 0.008 |
| **10** | 0.04 |
| **Naringinen** | 0.2 |
| **Novobiocin** | 0.004 |
| **Ciprofloxacin** | 0.3 |
